# Fast and accurate out-of-core PCA framework for large scale biobank data

**DOI:** 10.1101/2022.05.25.493261

**Authors:** Zilong Li, Jonas Meisner, Anders Albrechtsen

## Abstract

Principal Component Analysis (PCA) is widely utilized in statistics, machine learning, and genomics for dimensionality reduction and uncovering low-dimensional latent structure. To address the challenges posed by ever-growing data size, fast and memory-efficient PCA methods have gained prominence. In this paper, we propose a novel Randomized Singular Value Decomposition (RSVD) algorithm implemented in PCAone, featuring a window-based optimization scheme that enables accelerated convergence while improving the accuracy. Additionally, PCAone incorporates out-of-core and multithreaded implementations for the existing Implicitly Restarted Arnoldi Method (IRAM) and RSVD. Through comprehensive evaluations using multiple large-scale real-world datasets in different fields, we demonstrate the advantage of PCAone over existing methods. The new algorithm achieves significantly faster computation time while maintaining accuracy comparable to the slower IRAM method. Notably, our analyses of UK Biobank, comprising around 0.5 million individuals and 6.1 million common SNPs, demonstrate that PCAone accurately computes the top 40 principal components within a 9 hours. This analysis effectively captures population structure, signals of selection, structural variants, and low recombination regions, utilizing less than 20 GB of memory and 20 CPU threads. Furthermore, when applied to single-cell RNA sequencing data featuring 1.3 million cells, PCAone, accurately capturing the top 40 principal components in 49 minutes. This performance represents a 10-fold improvement over state-of-the-art tools.

## Introduction

Principal component analysis (PCA) is a popular approach that is often used to summarize the relationship between high dimensional observations. It has various applications, such as inferring population structure in genetics (Patterson et al. 2006) and clustering cells from single-cell sequencing experiments (Kiselev et al. 2019). The estimated principal components (PCs) are also widely used in downstream analyses. For instance, the PCs can be used as covariates in regression to account for confounding factors like population structure (Price et al. 2006). In addition, popular visualization methods, T-distributed Stochastic Neighbor Embedding (t-SNE) and Uniform Manifold Approximation and Projection (UMAP), are typically applied on the top PCs instead of the full dataset in order to reduce the computational burden (Diaz-Papkovich et al. 2019; Maaten et al. 2008). PCA is also employed as a general dimensionality reduction technique in both supervised and unsupervised settings by numerous existing statistics or machine learning models. However, performing a full-rank Singular Value Decomposition (SVD) is computationally infeasible when dealing with hundreds of thousands of observations (Tsuyuzaki et al. 2020; Privé, Luu, et al. 2020). To tackle this computational burden, fast PCA methods have been proposed to efficiently approximate the top PCs, avoiding the need for a full-rank SVD. These methods are widely used in both genetics and single-cell RNA sequencing (scRNA-seq) data analysis. These faster methods make trade-offs between accuracy, speed and memory usage, and they are typically employed to approximate a small number of the top principal components. Recently, Tsuyuzaki et al. 2020 conducted a systematic review of commonly used PCA algorithms and implemented various different algorithms. The most widely used fast PCA implementations are either based on the Implicitly Restarted Arnoldi Method (IRAM) (Lehoucq and Sorensen 1996), such as FlashPCA2 (Abraham et al. 2017), bigsnpr (Privé, Aschard, et al. 2018), and OnlinePCA.jl (Tsuyuzaki et al. 2020), or they are based on Randomized Singular Value Decomposition (RSVD) (Halko et al. 2011), such as FastPCA(Galinsky et al. 2016), TeraPCA (Bose et al. 2019) and OnlinePCA.jl (Tsuyuzaki et al. 2020). Other rapid methods also exist that can handle missing data such as PCAngsd (Meisner and Albrechtsen 2018), EMU (Meisner, Liu, et al. 2021) and ProPCA (Agrawal et al. 2020).

The Implicitly Restarted Arnoldi Method (IRAM), implemented first in the well-known ARPACK package (Lehoucq, Sorensen, and Yang 1998), serves as the backend for many packages in R (R Core Team n.d.), Python, and MATLAB. It is accurate but not pass-efficient, which has led to the recent popularity of Randomized Singular Value Decomposition (RSVD), despite its slightly reduced accuracy. However, the accuracy of RSVD can be improved by employing what are know as power iterations which essentially is an optimization step on the initial random matrix. While this improvement contributes to increased accuracy, it also introduces a higher number of passes over the data. As a result, this approach becomes computationally less efficient, particularly when dealing with data that exceeds available memory capacity. Effectively conducting PCA on extensive, high-dimensional dense data within the constraints of limited-memory computational environments continues to pose an ongoing challenge. Algorithms designed to operate out-of-core, without requiring all data to be stored in memory, offer a solution to the limited memory challenge. However, they can potentially be slower due to the incurred I/O operation costs associated with parsing data stored on disk or streamed (Yu et al. 2017; Abraham et al. 2017; Bose et al. 2019). Consequently, ensuring a pass-efficient implementation becomes crucial for out-of-core algorithms, aiming to minimize the frequency of data reads from the disk. Additionally, the capability for multithreading plays a vital role in enhancing software speed for both input data parsing and mathematical computations.

## Results

### Overview of PCAone framework

In this work, we present PCAone, an efficient and accurate out-of-core framework developed in C++. PCAone improves both the accuracy and the speed of the fast RSVD approach, while also including efficient implementations of existing PCA algorithms. The software is especially designed to efficiently process large-scale and high-dimensional data of different formats for biological research (Figure S3). Firstly, considering that RSVD is more pass-efficient but less accurate than IRAM, we propose and implement a new RSVD (Algorithm 2), referred to as PCAone. The method employs a window-based power iteration scheme on mini-batches of the input, that can achieve high accuracy within a few passes over the data (as illustrated in Figure S1 and S2). This feature renders the implementation well-suited for out-of-core computations, where I/O operations, such as reading data from the disk, are typically the limiting factor. Secondly, we implement an RSVD (Algorithm 1), referred to as PCAone*_H_*_+_*_Y_*, based on the approach described by Yu et al. 2017, but incorporating a power iteration scheme to improve accuracy. Notably, this approach utilizes a single pass over the data for each power iteration, constrasting with the two-pass strategy employed by other RSVD methods like OnlinePCA.jl*_Halko_* and FastPCA. This reduction in passes contributes to lower I/O costs. Furthermore, we acknowledge the importance of the IRAM algorithm as a suitable replacement for full SVD in estimating the top principal components (Privé, Aschard, et al. 2018; Tsuyuzaki et al. 2020). To assess the accuracy of alternative methods when full SVD is not computationally feasible, we introduce an efficient out-of-core implementation of the IRAM algorithm, referred to as PCAone*_Arnoldi_*. The implementation draws inspiration from FlashPCA2 (Abraham et al. 2017), enhanced with multithreading capabilities.

All three methods, PCAone, PCAone*_H_*_+_*_Y_* and PCAone*_Arnoldi_*, are implemented in both in-core and out-of-core modes, offering users the flexibility to optimize speed by utilizing available memory resources. However, in this paper, we concentrate exclusively on the out-of-core implementation, as it is the most suitable option for processing large-scale datasets. The details of PCAone algorithms and architecture are provided in Methods.

### Accuracy and iterations

We conducted a comprehensive comparative analysis of our three algorithms (PCAone, PCAone*_H_*_+_*_Y_*, PCAone*_Arnoldi_*) alongside various other software packages widely used for large-scale PCA in genetics. These packages include FlashPCA2, PLINK2 (a faster reimplementation of FastPCA, Chang et al. 2015), TeraPCA and ProPCA. To assess the overall accuracy of all estimated principal components (PCs), we computed the mean explained variance (MEV) between the estimated PCs and those obtained from the full-rank SVD.

First, we analyzed a dataset of four genetically similar East Asian populations sourced from the 1000 Genomes Project. These populations consist of two Han Chinese population (CHB, CHS), a Dai Chinese (CDX) and a Vietnamese population (KHV), each comprising approximately 100 individuals. The dataset included 5,675,746 common Single Nucleotide Polymorphisms (SNPs) shared among those populations. Full-rank SVD can be performed to obtain the true PCs for this dataset due to its small sample size. Given the genetic similarity among the four East Asian populations and the gradual decay of eigenvalues in the dataset (Figure S5), a higher number of iterations is expected to achieve high accuracy (Halko et al. 2011; Yu et al. 2017). We define epochs as the number of times the data is read from the disk for the out-of-core methods, and as the number of iterations for in-core methods like PLINK2 (FastPCA) and ProPCA. The results for different top *K* PCs are shown in Figure 1A. Notably, our proposed PCAone algorithm consistently uses 7 epochs for all choices of *K*. In contrast, the other algorithms use more epochs for higher *K*, which is determined by both the algorithms convergence efficiency and their stopping criteria (Table S5). For *K* = 2, most methods produce nearly identical top 2 PCs compared to the full-rank SVD, with the exception of PLINK2 (FastPCA) and ProPCA. These two methods fail to accurately capture the difference between the Dai and Vietnamese (Figure 1B). The main cause for their low accuracy at *K* = 2 is their stopping criteria, which is solely based on the choice of *K*. With higher choices of *K*, their accuracy improves. Regarding memory consumption, most methods exhibited small memory requirements for this dataset, demonstrating near constant usage across different *K* settings. However, PLINK2 (FastPCA) showed increasement of memory consumption for high *K* value (Figure 1C). The comparison of accuracy across different choices of *K* (Figure 1D) revealed that PCAone and the two IRAM methods (PCAone*_Arnoldi_*, FlashPCA2) consistently achieved very high accuracy, while the other methods performed worse for some choices of *K*. The accuracy remained consistent when performing the analysis on random subsets of one million SNPs (Figure S4), indicating minimal uncertainty in the MEV estimates. Disparities in accuracy among the RSVD methods are primarily explained by their stopping criteria and thus the number of epochs they use. The notable exception is PCAone which converges faster per epoch than the conventional RSVD method with standard power iteration (Figure 1E). The faster convergence of PCAone is achieved by performing many power iterations each epoch on subsets of the data (Figure S2).

**Figure 1:**
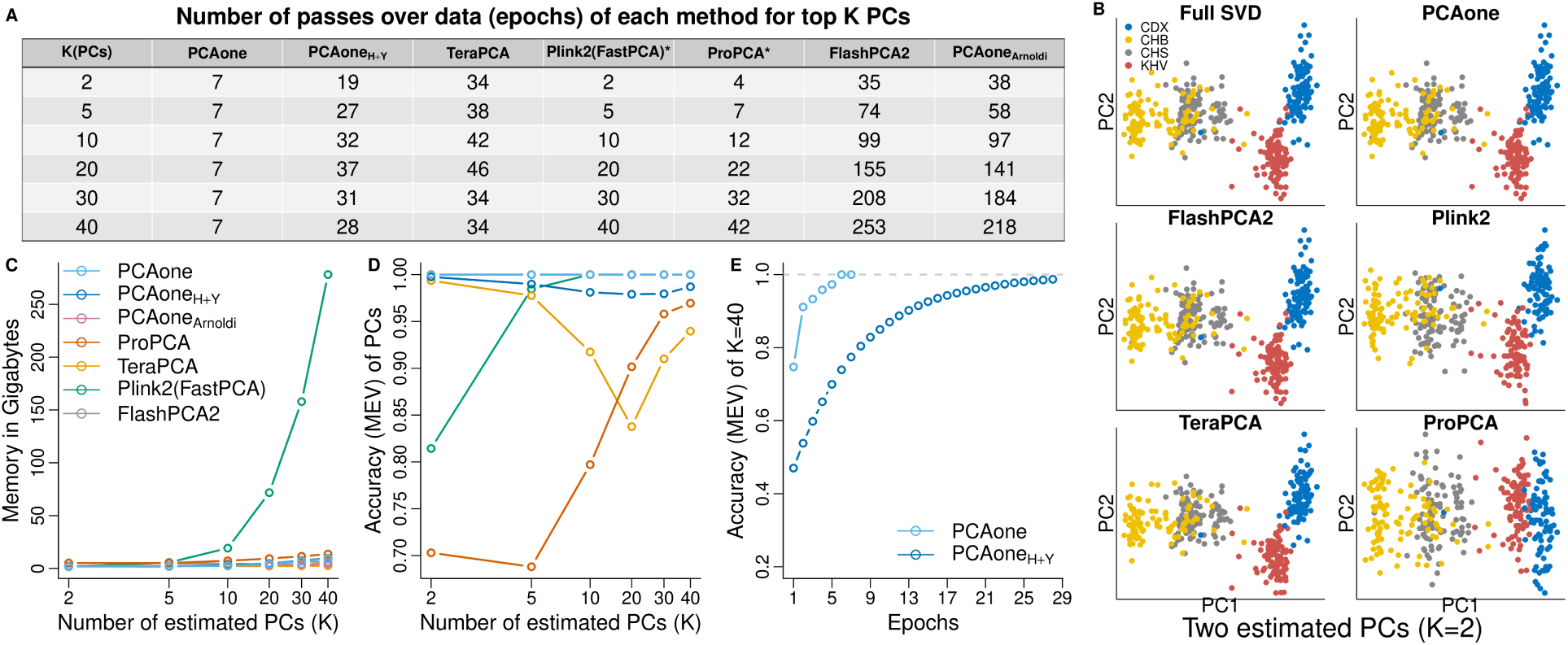
Performance on the East Asian data from 1000 Genomes Project. PCA performance of different software with various number of inferred PCs (K) based on 2,000,000 SNPs and 400 individuals from four East Asian populations. **(A)**, Number of epochs used by each software. * Not out-of-core, only allows for in-core computation with this number of iterations. **(B)**, Two estimated PCs *K* = 2 for each methods including the true full SVD . **(C)**, Memory usage as a function of *K* (log scale at x-axis). **(D)**, Accuracy (MEV) compared to the true full-rank SVD as a function of *K* (log scale at x-axis). **(E)**, Convergence of PCAone and PCAone*_H_*_+_*_Y_* shown as accuracy per epoch.

We extended our analysis to encompass all populations from the 1000 Genomes Project (Figure S7) and the Human Genome Diversity Project (Figure S8), where a more pronounced population structure exists, leading to a larger portion of the variation being explained by the top PCs (Figure S5). Both datasets contain individuals from diverse populations, including admixed individuals. Across various selected numbers of PCs, most methods achieved high accuracy (MEV > 0.99), and PCAone consistently required only 7 or fewer epochs to achieve very high accuracy. The number of epochs needed by PCAone and PCAone*_H_*_+_*_Y_* depends on the estimated number of PCs, the number of individuals, and the number of sites, as explored in Figure S12 and S13. Importantly, PCAone has high accuracy of MEV > 0.99 in all analyses, and in most cases, it required only 7 epochs. However, there are instances with relative smaller datasets and a high number of estimated PCs where it may require 8 or 9 epochs for optimal performance. In contrast, the conventional RSVD methods, PCAone*_H_*_+_*_Y_*, experiences a more significant increase in the number of epochs needed, requiring up to 24 epochs.

### Runtime and memory usage

We assessed the scalability and performance of PCAone and the other methods, using random subsets extracted from the UK Biobank genetic data. We randomly sampled subsets of individuals and SNPs that are small enough for most of the methods to run. It’s noteworthy that all methods executed until convergence, governed by their respective stopping criteria (Table S5). Across these datasets, all methods exhibited high accuracy (MEV > 0.999, Table S1) and low minimum sum of squared error (minSSE, Table S2). We further compared the wall-clock time and memory consumption for calculating the top *K* = 40 PCs. As shown in Figure 2, all methods exhibited a linear relationship between speed and number of individuals or SNPs. The memory usage remained near constant for the out-of-core implementations (PCAone, PCAone*_H_*_+_*_Y_*, PCAone*_Arnoldi_*, FlashPCA2, TeraPCA), whereas it exhibited a linear relationship for ProPCA and PLINK2. When using 20 threads, our IRAM implementation, PCAone*_Arnoldi_*, outperformed FlashPCA2 by a factor of 10*×* for the largest dataset. This performance gain is primarily attributed to the multithreading capability of PCAone*_Arnoldi_*, which enables parallel processing of input files and calculations. Likewise, our out-of-core RSVD implementations, PCAone and PCAone*_H_*_+_*_Y_*, are much faster than the other methods including the in-core methods of PLINK2 (FastPCA) and ProPCA, which do not read in data from the disk during each iteration. Throughout all benchmarking, PCAone is on average 2*×*, 5*×*, 9*×*, 12*×* and 30*×* faster than PCAone*_H_*_+_*_Y_*, PLINK2, ProPCA, TeraPCA and FlashPCA2, respectively, while utilizing little memory. The main advantage of PCAone is that it converges in less epochs compared to other methods such as TeraPCA, PLINK2 and ProPCA, given that they share similar time complexity per epoch (Table S6). Moreover, PCAone shows a more efficient multithreaded implementation, which is also used when reading data from disk. This is demonstrated through a benchmarking analysis of PCAone for analyzing genotype dosages in BGEN format, as compared to genotype calls in PLINK format (Table S3). Notably, the BGEN format contains float values between 0 and 1, posing a greater complexity in storage and parsing compared to the binary PLINK format. As shown in Table S3, PCAone requires only three times longer for processing genotype dosages.

**Figure 2:**
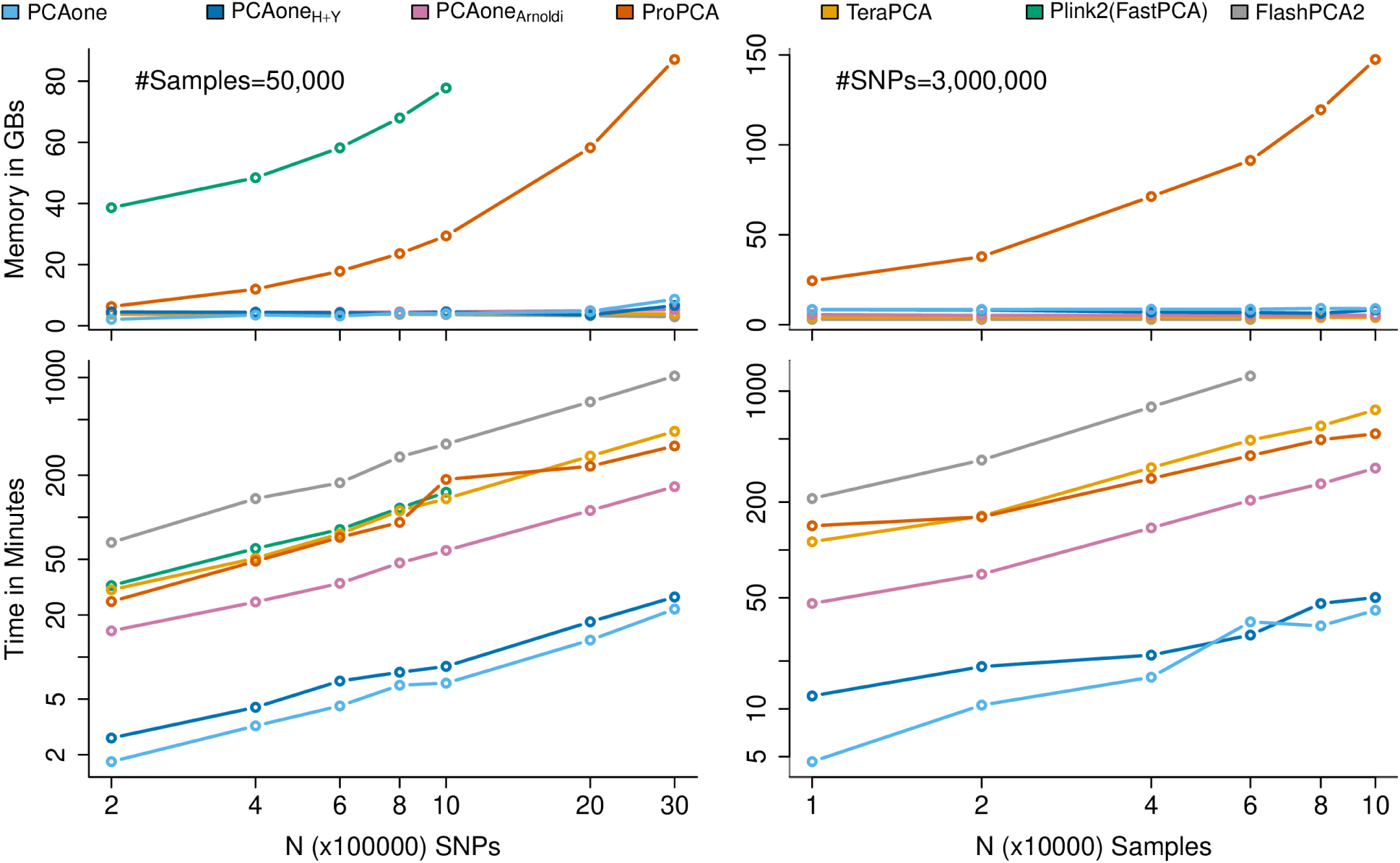
Runtime and memory usage. Runtime and memory usage for calculating top K=40 PCs in either random subsets of common SNPs (left column) for 50,000 individuals or random subsets of individuals (right column) for 3,000,000 common SNPs. We used 20 CPU threads for all programs except FlashPCA2 which does not support multi-threading. Detailed commands are included in Benchmarking section. PLINK2 (FastPCA) ran out of memory when using more than 1,000,000 SNPs.

We conducted further analysis on a dataset comprising 487,409 individuals and 6,133,304 common SNPs sourced from the UK Biobank. It was not feasible to run ProPCA and PLINK2 (Fast-PCA) as they needed more than 180 GBs of allocated memory. The accuracy and resource usage are summarized in Table 1. Among the methods, only PCAone and PCAone*_H_*_+_*_Y_* successfully completed the task within a day, with runtimes of 9.2 hours and 12.6 hours, respectively. This runtime also includes the time for permuting data as required by PCAone, which took approximately one hour. One the contrary, TeraPCA required 49 hours, PCAone*_Arnoldi_* took 102 hours to complete, and FlashPCA2 was not expected to finish within a month. Again, we calculated accuracy using both MEV and minSSE for the three RSVD implementations compared to PCs estimated by the highly accurate IRAM method in PCAone*_Arnoldi_*, which showed PCAone*_H_*_+_*_Y_* and TeraPCA had slightly lower MEV and higher minSSE compared to PCAone. Moreover, to understand the impact of the marginally lower accuracy on the PCs, we also ran PCAone*_H_*_+_*_Y_* with a stricter stopping criterion, exploring the number of epochs required to achieve comparable accuracy to PCAone. We further illustrated this effect by presenting the minSSE per PC and highlighting the individual points with the least accuracy for each method (Figure S10 and S11). These visualizations illustrate that PCAone, when employing default stopping criteria, yields outcomes akin to those of the IRAM method.

**Table 1:** Performance on UK Biobank imputed data with 487,409 individuals and 6,133,304 SNPs for estimating *K* = 40 PCs using 20 CPU threads. Time is given in hours and RAM is in gigabytes. *^α^* refers to I/O time for permuting data. Wall time includes I/O time and computation time. PCs estimated by PCAone*_Arnoldi_* are used as the true value for calculating accuracy (MEV and minSSE). PCAone*_H_*_+_*_Y_* * using a stopping criteria to achieve similar accuracy as PCAone. Accuracy of individual PC is given in Figure S7 and S8.

| Program | Wall Time(h) | IO Time(h) | Epochs | RAM(GB) | MEV | minSSE |
| --- | --- | --- | --- | --- | --- | --- |
| PCAone | 9.2 | $4.9+0.9^{\alpha}$ | 7 | 18.64 | 0.999996 | 0.000362 |
| $\text{PCAone}_{H+Y}$ | 12.6 | 6.2 | 10 | 17.09 | 0.999664 | 0.046124 |
| $\text{PCAone}_{H+Y}^*$ | 23.3 | 16.1 | 17 | 17.11 | 0.999997 | 0.000223 |
| TeraPCA | 49.3 | 38.8 | 20 | 9.40 | 0.999432 | 0.048754 |
| $\text{PCAone}_{Arnoldi}$ | 102.7 | 85.8 | 124 | 23.27 | - | - |

PLINK2 (FastPCA) and ProPCA couldn't run in limited memory ( $>180$ GBs).
FlashPCA2 couldn't finish in limited time ( $>30$ Days).

### Analyses on UK Biobank data

To underscore the practical utility of conducting PCA on extensive genetic datasets, we performed PCA of the UK Biobank imputed data, encompassing 487,409 individuals and 6,133,304 SNPs. Im-portantly, we refrained from performing Linkage Disequilibrium (LD) pruning in order to capture both local and global genetic patterns. A closer examination of the distribution of SNP loadings for each Principal Component (PC) provided insights into their significance (see Figure S6). PC1, PC2, and PC4 effectively captured the population structure, as evident in Figure 3B and Figure 3C. Based on previous studies, we annotated the SNPs with the highest loadings for each PC(Supplemental Table S1). The SNP *rs16891982* had the highest loadings in both PC1 and PC2 (as shown in Figure 3A) and also ranked as the third highest loading in PC4. This SNP is situated within the skin pigmentation gene *SLC45A2*, which is known to be under positive selection (Beleza et al. 2013). Thus, this observation suggests the feasibility of implementing a PC-based selection method (Galinsky et al. 2016). However, as shown in Figure 3D, the majority of PCs tend to capture regions of low recombination, such as centromeres and inversions. Other interesting peaks include the large inversion found on Chromosome 8 (Porubsky et al. 2022; Bansal et al. 2007) and the HLA region on Chromosome 6.

**Figure 3:**
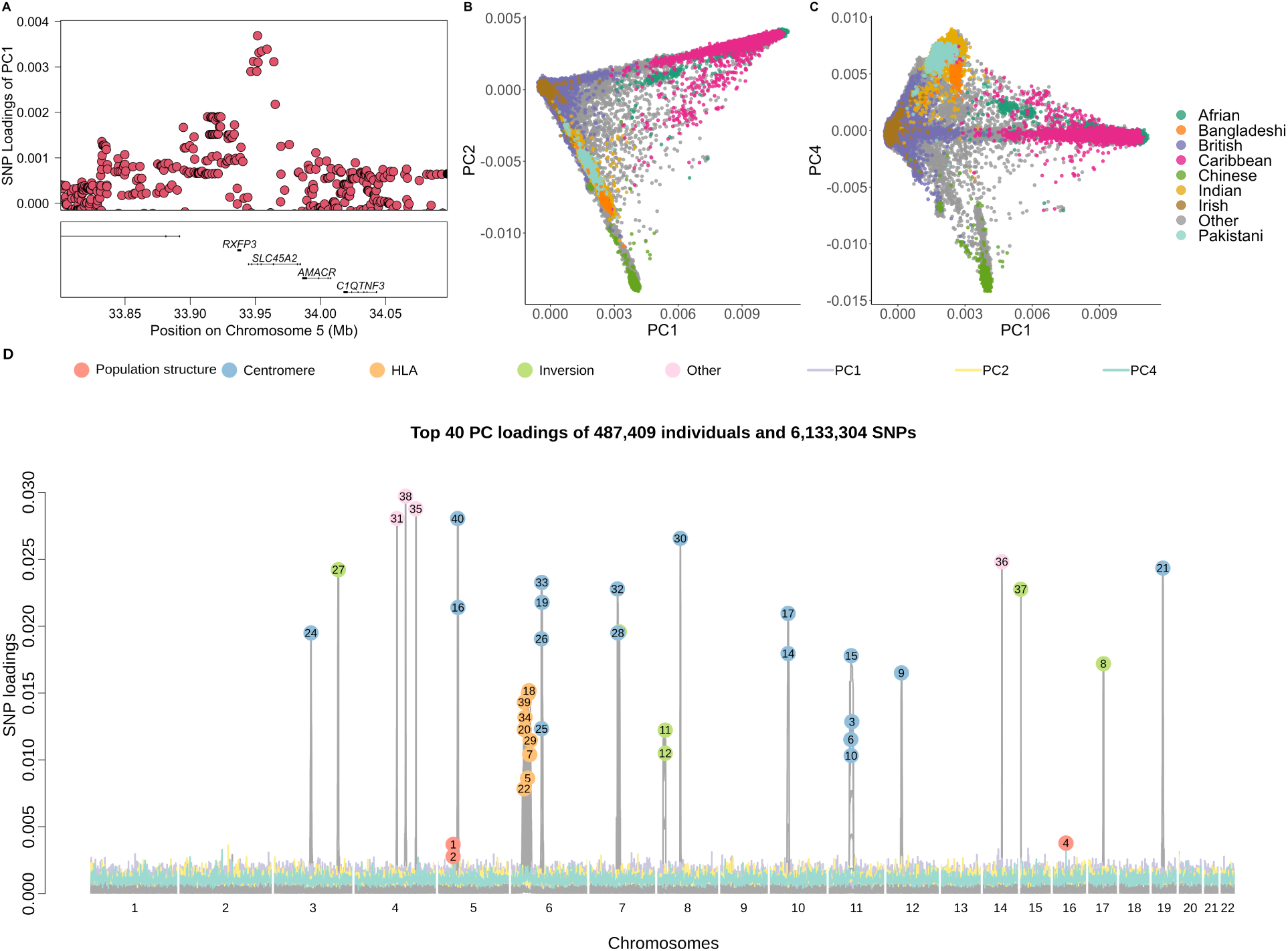
PCA of the whole UK Biobank imputed dataset. **(A)**, LocusZoom of variant rs16891982 with highest loadings for PC1 and PC2. **(B)** and **(C)**, Population structure showed in PC1 vs PC2 and PC1 vs PC4 for all 487,409 individuals, colored by country or region label. Other labels are merged into the category “other”. **(D)**, SNP loadings for the top 40 PCs. Peaks are annotated and colored by the type of genomic region they are located in.

An alternative approach to conducting PCA on imputed genotype calls involves accounting for genotype uncertainty by analyzing genotype dosages. These dosages are represented as floating-point numbers in the BGEN format and require more storage space, and a difference manifested in PCAone being three times slower (as detailed in Table S3). The correlation between Principal Components (PCs) obtained from imputed genotypes and those derived from dosages is notably high, as shown in Figure S9. Consequently, for this specific dataset, the choice between imputed genotypes and dosages has minimal impact. However, when analyzing only the genotype calls from the LD-pruned UK Biobank SNP chip data, a noteworthy distinction emerges. In this scenario, only PC1, PC2, and PC4, which capture the population structure, have high correlation between the datasets. The data is small enough to perform full SVD which has very high MEV of 0.9999385 and 0.9999006 with PCAone and PCAone*_H_*_+_*_Y_* respectively.

### Analyses on single cell and bulk RNA sequencing data

Within the PCAone software, we have implemented a range of PCA algorithms and extend support to diverse data types. To demonstrate the application of PCAone in the relm of single cell RNA sequencing (scRNA-seq), we performed PCA using one of the largest scRNA-seq datasets available. This dataset comprises 1,306,127 mouse brain cells and 23,771 genes. Notably, the calculation of the top 40 PCs took 71 minutes and 53 minutes for PCAone and PCAone*_H_*_+_*_Y_* respectively. We compared the performance with OnlinePCA.jl*_Halko_*, which was recently reported as the fastest out-of-core implementation available for scRNA-seq (Tsuyuzaki et al. 2020). By default, PCAone assumes the data has been normalized by the users but it also supports the common normalization method for scRNA-seq (see Methods). Notably, OnlinePCA.jl separates the normalization and PCA calculation into two processes, whereas PCAone performs file parsing, normalization and PCA simultaneously. We evaluated the time for I/O, normalization and PCA computation separately for each program. As Table 2 shown, PCAone consumed less I/O and computing time than

**Table 2:** Performance on scRNA-seq data with 1,306,127 cells and 23,771 genes for estimating the top K=40 PCs. Time is given in minutes and RAM in gigabytes. Wall time includes IO time. * normalization of data is performed in a separate step by OnlinePCA.jl while PCAone did this on the fly. For wall time, we have not included the time needed by OnlinePCA.jl*_Halko_* to make their binary format used for quick I/O or the time of permutation potentially needed for PCAone. PCs estimated by PCAone*_Arnoldi_* are used as the true value for calculating accuracy (MEV and minSSE). All programs used 20 CPU threads.

| Program | Wall Time(m) | IO Time(m) | Epochs | RAM(GB) | MEV | minSSE |
| --- | --- | --- | --- | --- | --- | --- |
| PCAone | 49 | 31 | 10 | 8.97 | 0.999957 | 0.004171 |
| PCAone<sub>H+Y</sub> | 42 | 30 | 8 | 6.82 | 0.999925 | 0.007328 |
| OnlinePCA.jl<sub>Halko</sub> | 461+100* | 300 | 8 | 1.71 | 0.954103 | 7.102932 |
| PCAone<sub>Arnoldi</sub> | 484 | 469 | 103 | 5.73 | - | - |

OnlinePCA.jl leveraging 20 threads. Despite using CSV format as input, PCAone demonstrated over 5*×* faster performance than OnlinePCA.jl using 20 threads. Given the impracticability of performing full SVD on such a large dataset, we derived MEV and minSSE values using the results from the IRAM method (PCAone*_Arnoldi_*). To underscore that PCAone*_Arnoldi_* is a suitable substitute for full SVD, we also performed the analysis on a subset of 12000 cells, where the accuracy remains the same when replacing full SVD with PCAone*_Arnoldi_* (Table S4). In Figure 4, a comparative analysis of the top 40 PCs showcased the lower accuracy of OnlinePCA.jl*_Halko_* for tailing PCs. Conversely, both PCAone and PCAone*_H_*_+_*_Y_* consistently achieved high accuracy. Notably, the PCA outcomes from PCAone exhibited visual similarity to the slower IRAM method PCAone*_Arnoldi_* unlike OnlinePCA.jl*_Halko_*. We also analyzed bulk RNA-seq data from the GTEx project, involving 56,200 gene transcripts across 54 tissues from 943 individuals (Figure S15). Here, both PCAone and PCAone*_H_*_+_*_Y_* converged fast and had high accuracy, even when estimating 100 PCs. It’s noteworthy that, in contrast to the genetic data, PCAone*_H_*_+_*_Y_* converged with slightly fewer epochs compared to the 7 required by PCAone.

**Figure 4:**
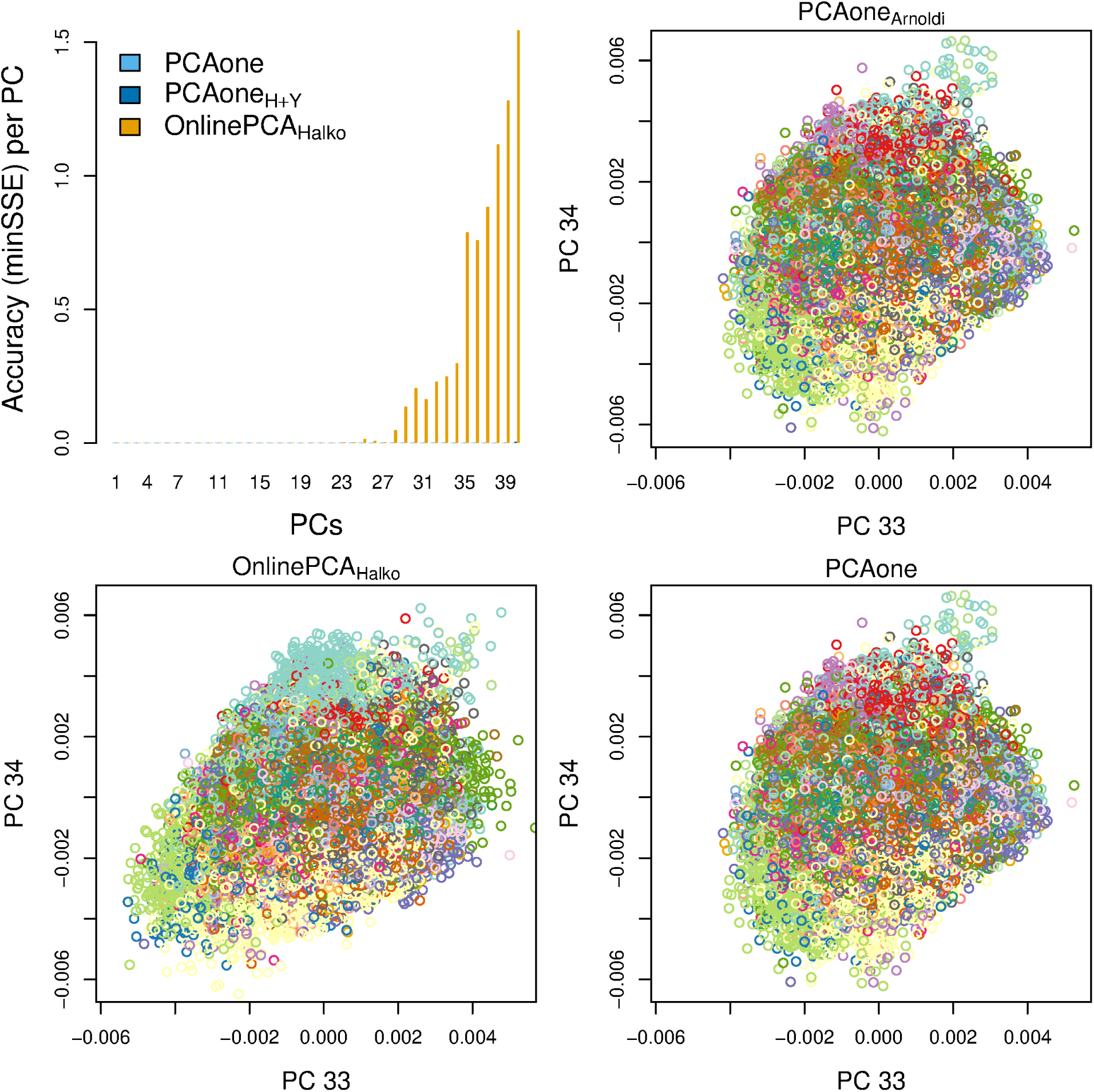
Accuracy of individual PCs for scRNA-seq data with 1,306,127 cells and 23,771 genes. The top left plot is the accuracy (minSSE) of each PC where results by PCAone*_Arnoldi_* are used as the true value. PC 33 and 34 are shown for PCAone and OnlinePCA.jl*_Halko_* to illustrate the difference compared to the accurate PCAone*_Arnoldi_*.

## Discussion

We presented PCAone, an out-of-core PCA framework tailored for large-scale data analyses. Through various datasets, we have demonstrated the advantage of our implemented IRAM method, PCAone*_Arnoldi_*, which is 10*×* faster than the commonly used IRAM method FlashPCA2. The speed advantage is primarily attributed to its multithreading capability. However, the IRAM method (PCAone*_Arnoldi_*) can be significantly slower than the RSVD methods (PCAone*_H_*_+_*_Y_* and PCAone) in terms of the number of iterations required for convergence. Throughout our benchmarking, our novel window-based RSVD algorithm, PCAone, consistently showed high accuracy and meaningful speed improvements over the existing RSVD algorithms. This is particularly evident when the eigenvalues of the estimated principal components (PCs) decay slowly, as PCAone requires significantly fewer epochs to converge, making it substantially more efficient than conventional RSVD methods. We have demonstrated this point using genotype data sourced from the genetically similar East Asian populations within the 1000 Genomes Project, where PCAone converges in only 7 epochs in contrast to the other methods that needed many times more epochs. In addition to its accelerated speed, PCAone also performs nearly as well as the IRAM methods (PCAone*_Arnoldi_* and FlashPCA2) and produces visually indistinguishable PCs. In contrast, other RSVD-based algorithms performed worse for some datasets such as the East Asian dataset. The accuracy of PLINK2 (FastPCA) and ProPCA increased when estimating a higher number of PCs, which is expected as the number of optimization steps is determined by the number of PCs to be estimated. Most of the methods allow users to define their own stopping criteria, which can be used to increase accuracy at the expense of speed. PCAone typically converges in exactly 7 epochs using the default setting of 64 mini-batches for its expanding window optimization scheme. By performing optimization on subsets of the data, the algorithm approximates the solution derived from the full dataset. And the optimization steps of PCAone utilize expanding window sizes, with epoch 6 and 7 being the first two consecutive iterations that involve the entire dataset. Consequently, PCAone achieves convergence in fewer epochs than other RSVD methods due to its multiple updates of the initial random matrix Ω within the first 5 epochs. The window-based optimization used in PCAone is essentially an incremental learning approach, commonly referred to as mini-batch or online learning. This technique finds widespread use in statistics and machine learning for modeling large-scale datasets (Neal et al. 1998; Cappé et al. 2009; Kingma et al. 2017; Kuhn et al. 2020). Other methods like Sequential Karhunen–Loeve (SKL) transform algorithm (Levey et al. 2000; Ross et al. 2008) can be viewed as an incremental PCA method. While it has been shown that incremental learning can speed up the convergence of a stochastic model, it can lead to worse performance depending on the nature of the data and settings of the algorithm. For instance, there is a “forgetting factor” parameter in SKL that determines how much weight the algorithm should put on a new mini-batch of data, which can have a big impact on performance. In contrast, to achieve the best of both worlds, we incrementally expand the mini-batch data in the first 5 epochs until the window encompasses the entire dataset, at which point the algorithm aligns with conventional power iterations. Thus, the algorithm shares the same theoretical foundation as the conventional RSVD by Halko et al. 2011. We realize that it is challenging to determine the optimal number of power iterations to use for RSVD. In general, RSVD packages just use 3 or 10 power iterations based on experiences, which may be not optimal depending on the dataset. For instance, PCAone*_H_*_+_*_Y_* used slightly fewer epochs than PCAone for the RNA-seq data, which may indicate there is no need to do many power iterations for those datasets. Nevertheless, the advantage of the PCAone algorithm is that it performs many power iterations in each pass through the data with efficient multithreading. However, a disadvantage of this expanding window scheme is that performing calculations on multiple machines is less straightforward than PCAone*_H_*_+_*_Y_*, which can be easily adapted into a distributed PCA method (Elgamal et al. 2015), with each machine handling calculations for separate blocks of data.

Beyond the three fundamental PCA algorithms, one can take advantage of the PCAone frame-work to implement other methods that build upon the PCA algorithms. For instance, methods involving iterative PCA computations can greatly benefit from the PCAone algorithm. In this context, we implemented EMU (Meisner, Liu, et al. 2021) and PCAngsd (Meisner and Albrechtsen 2018) methods within PCAone to infer population structure for genetic data that exhibit missingness and uncertainty. The PCAone framework is highly optimized for out-of-core computations and is implemented as a multithreaded C++ software. Another way to solve big memory issue is using memory-mapped file such as bigsnpr (Privé, Aschard, et al. 2018). Additionally, we have implemented the PCAone algorithm as an R package, allowing users to seamlessly incorporate PCAone into the R ecosystem (R Core Team n.d.). This includes the ability to work with memory-mapped objects using packages like bigmemory (Kane et al. 2013).

## Methods

### PCAone algorithms

Our proposed algorithm is based on the randomized SVD introduced by Halko et al. 2011. We briefly explain the underlying idea of this algorithm, which aims to find a near-optimal orthonormal basis *Q^m×k^* for input matrix *A^m×n^* such that *A ≈ QQ^T^ A* with *k « m, n*. Here *n* denotes the number of samples and *m* denotes the number of features. With such a *Q*, the right eigenvectors of *A* are approximated by the right eigenvectors of the low dimensional *B* = *Q^T^ A*. The RSVD algorithm involves multiplying the input *A* with a random low-dimensional matrix Ω*^n×^* drawn from a standard normal distribution. Then QR factorization is performed on the resulting product (*Q, R* = *qr*(*A*Ω)). The entire procedure usually goes though the data twice. Recently, Yu et al. 2017 proposed a more efficient version of this algorithm that goes through the input matrix only once and includes re-orthogonalization of Q to alleviate round of errors. However, these single-pass algorithms are not always very accurate. Halko et al. 2011 also proposed a more accurate algorithm, which involves optimizing Ω through another orthogonalization process (Ω*, R* = *qr*(*A^T^ Q*)). These two steps are the so-called power iteration steps. As noted by Yu et al. 2017 it is possible to incorporate the power iteration scheme into their algorithm. We have implemented such an algorithm in Algorithm 1. It is important to note that when we want to estimate *k* PCs we will use a matrix of *Q* with *l* = *k* + 10 columns. This padding of 10 extra PCs helps alleviate the issues of eigenvectors order switching in the optimization step. Algorithm 1 allows us to read the data in small batches once per iteration, making it very useful for large datasets that does not fit in memory. By adding power iterations to it, we can achieve higher accuracy.

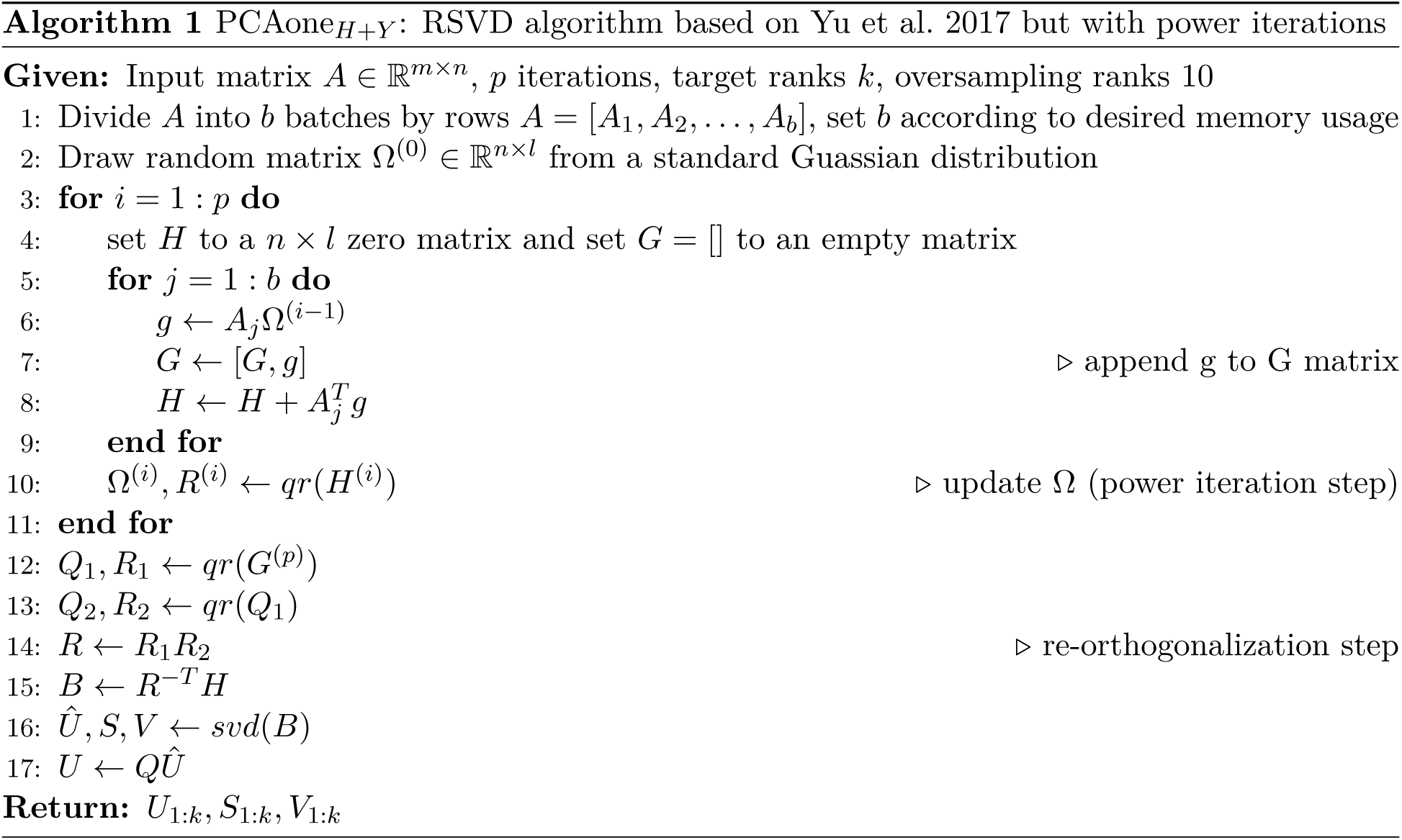

However, Algorithm 1 is not ideal for scenarios where many power iterations are needed to achieve high accuracy. Hence, we propose Algorithm 2 to speed up its convergence and achieve high accuracy within fewer passes through the data by incorporating multiple updates of Ω (power iterations) in each epoch. We denote the number of times the data must be read from the disk as epochs. Firstly, we permute the input matrix *A* by rows (features) and divide it into 64 blocks of approximately equal size. This ensures each block of the data is a random subset of the features, which is important for datasets where the adjacent features are correlated. For instance, the genetic variants are stored based on its position and exhibit local correlation due to linkage disequilibrium. In the first epoch, we start with a window size of 2 blocks of data and utilize this subset of data to perform the power iterations to update Ω. Subsequently, we slide the window by one block (step size=1) so that the next iteration, which updates Ω, is performed on block 2 and 3. After sliding through the whole data (one epoch), we double both the step size and the window size until the 6th epoch, where the window size equals the entire input matrix as illustrated in Figure S2. Thereafter, the power iterations are performed in the same way as in the conventional RSVD. The algorithm terminates when the difference of eigenvectors between two successive epochs is smaller than an user defined threshold (default 1 *− MEV <* 10*^−^*^4^). This often occurs after 6 epochs, where a second update of Ω based on the entire data leads to only a very small change of Ω and thus the estimated PCs. For simplicity, the PCAone algorithm shown in Algorithm 2 is presented as if the whole block of data is read into memory. However for large datasets, we only keep small batches of the data matrix *A* in memory as shown in Algorithm 1. Additionally, the sign of each of its columns will change in each iteration due to the QR factorization-based updates of Ω. We therefore ensure consistency between iterations by flipping the sign if necessary.

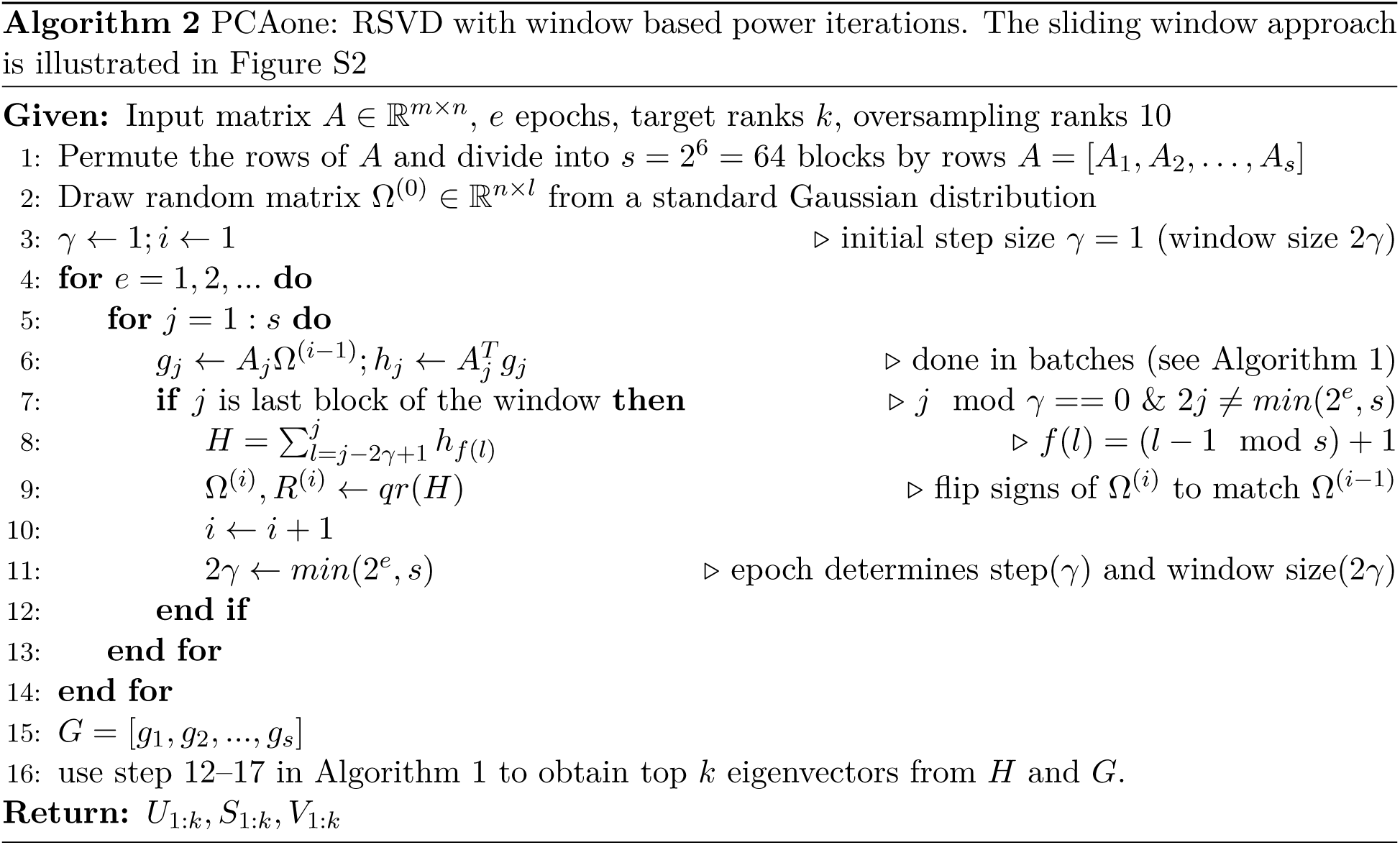

The major difference between PCAone*_H_*_+_*_Y_* (Algorithm 1) and PCAone (Algorithm 2) lies in the number of power iterations performed for each epoch. In the first five epochs, PCAone updates Ω 124 times while PCAone*_H_*_+_*_Y_* only performs 5 updates. Starting from the 6th epoch, the two algorithms are the same and Ω is updated only once per epoch. The theoretical foundation for both is the same as demonstrated by Halko et al. 2011.

### Convergence and accuracy measurement

To measure the similarity of two eigenvector matrices, we employ the mean explained variance (MEV) (Galinsky et al. 2016; Agrawal et al. 2020) and minSSE. Both measures are insensitive to the order of the eigenvector matrix. Assuming we have a true set of PCs (*v*_1_*, v*_2_*, …, v_k_*), we evaluate the accuracy of a set of estimated PCs (*u*_1_*, u*_2_*, …, u_k_*) as follows:

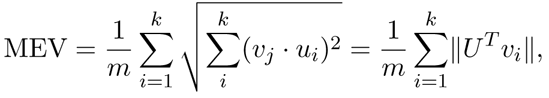

where *U* is a matrix with the estimated PCs as column vectors.

MEV, representing the overall accuracy, can become very small for large matrices even if there are substantial differences between some values. Therefore, we also employ a metric that considers column reordering, relying on the sum of squared error between the two matrices. The metric is referred to as minSSE and it is defined as

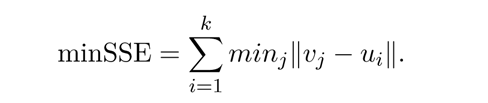

The minSSE is a measure of the minimum squared distance between matrix U and the columns of matrix V, computed based on a combination of *v_j_*.

In certain RSVD implementations, users are required to manually specify the number of power iterations, leading to a lack of a clear and explicit accuracy metric. For PCAone*_H_*_+_*_Y_* and PCAone, we can obtain the eigenvectors and eigenvalues after each power iteration, such that we can easily compare two sets of estimated PCs from successive power iterations and implement an explicit stopping criteria. As shown in Figure S14, our stopping criteria based on 1 *−* MEV *≤* 10*^−^*^4^ between successive epochs work well for achieving high accuracy.

### Dataset

To benchmark the performance of all PCA methods, we use multiple real-world datasets.

#### East Asian data

We used genotype data from the high depth whole genome sequencing data of all 400 individuals in four genetically akin East Asian populations from 1000 Genomes Project(https://www.internationalgenome.org/data-portal/data-collection/grch38): two Han Chinese populations (CHB, CHS), a Dai Chinese (CDX) population and a Vietnamese (KHV) population. We analyzed the 5,675,746 common SNPs with minor allele frequency (MAF) greater than 0.05 among these individuals. To quantify the uncertainty of the MEV estimates, we performed the analyses on ten different random subsets of one million SNPs.

#### HGDP and 1000 Genomes data

We analyzed the genotype data from all populations in the Human Genome Diversity Project (HGDP) and the 1000 Genomes Project (https://www.internationalgenome.org/data-portal/ data-collection/grch38), which are more diverse than the East Asian data. The HGDP data was downloaded from the latest release (v3) of gnomAD (https://gnomad.broadinstitute.org/ downloads#v3-hgdp-1kg). For both datasets, we only kept SNPs with MAF > 0.05 for the analyses (Figure S7, S8). To benchmark how PCAone scales with the number of features, we created multiple datasets by randomly sampling the common SNPs from the 1000 Genomes data in Figure S12.

#### UK Biobank data

The UK Biobank data is granted under Application Number 32683 from the UK Biobank Resource https://biobank.ctsu.ox.ac.uk/crystal/label.cgi?id=100319. We analyzed all 6,133,304 common SNPs with MAF > 0.05 for all individuals in the UK Biobank imputed dataset. For benchmarking the speed, memory usage and scalability of different methods, we created multiple datasets by randomly sampling individuals and common SNPs from this dataset. To show the versatility of PCAone, we also analyzed the genotype dosage in BGEN format for the same imputed SNPs in Figure S9. In addition to the imputed data, we analyzed the UK Biobank Axiom™ Array data with 498,444 SNPs after LD-pruning by *plink --indep-pairwise 50 10 0.5* in Figure S9.

#### Single cell and bulk RNA sequencing data

Additional to genetic data, we benchmarked PCAone’s performance on the public single cell RNA sequencing (scRNA-seq) data with raw gene expression counts from https://support.10xgenomics.com/single-cell-gene-expression/datasets/1.3.0/1M_neurons. The data consists of 1,306,127 mice brain cells from 60 clusters and and 23,771 genes generated with the 10x Chromium (Zheng et al. 2017). The label of each cell belonging to which cluster was used to color the cell point in PCA plots. We also randomly sampled 200 cells within each cluster (12000 cells in total) and 23,771 genes, from which we can calculate the full-rank SVD to validate that we can use our IRAM implementaion PCAone*_Arnoldi_* to measure accuracy.

Another common type of count data is the bulk RNA sequencing data. We analyzed the GTEx bulk RNA-seq data (https://storage.googleapis.com/gtex_analysis_v8/rna_seq_data/GTEx_ Analysis_2017-06-05_v8_RNASeQCv1.1.9_gene_tpm.gct.gz) in Figure S13, which consists of 56,200 gene transcripts counts (TPM) and 17,382 samples from 54 tissues and 943 individuals (https://raw.githubusercontent.com/broadinstitute/gtex-v8/master/data/GTEx_Analysis_ v8_RNAseq_samples.txt). For normalization of the bulk RNA-seq data, we used log transformation which is supported in PCAone by using the *--scale 1* option.

### Benchmarking

For benchmarking on genetic data in Figure 2, we used 20 threads for all programs except for FlashPCA2 which does not support multithreading. Specifically, we used PCAone(v0.3.5), PLINK2(v2.00a2.3LM), ProPCA(commit e94c972), TeraPCA(commit b4f8293) and FlashPCA2(v2.1) with the following settings:

- PCAone*_Arnoldi_*: --svd 0 --memory 4 -n 20 -k 40
- PCAone*_H_*_+_*_Y_* : --svd 1 --memory 4 -n 20 -k 40
- PCAone: --svd 2 --memory 4 -n 20 -k 40
- PLINK2: --pca approx 40 --threads 20
- ProPCA: -vn -cl 0.001 -a -nt 20 -k 40
- TeraPCA: -filewrite 1 -print 2 -memory 4 -n 20 -nsv 40
- FlashPCA2: --memory 4100 --verbose -d 40

For benchmarking on scRNA-seq data Table 2, we also used 20 threads for PCAone(v0.3.5) and OnlinePCA.jl(v1.0.5) with the following settings.

- PCAone*_Arnoldi_*: --svd 0 --scale 2 --memory 4 -n 20 -k 40
- PCAone*_H_*_+_*_Y_* : --svd 1 --scale 2 --memory 4 -n 20 -k 40
- PCAone: --svd 2 --scale 2 --memory 4 -n 20 -k 40
- OnlinePCA.jl*_Halko_*: –niter 3 –dim 40 –scale log –threads 20

For more detailed information, we provide the reproducible workflow on our GitHub repository https://github.com/Zilong-Li/Papers/tree/main/pcaone. All benchmarking were performed on a machine with 192 GB of memory and 88 CPU threads available (CPU model: Intel(R) Xeon(R) Gold 6230 CPU @ 2.10GHz).

### Normalization for genetic data

For inferring population structure with genetic data, PCAone supports genotypes (PLINK binary format) or genotype dosage (BGEN format) as input. Let *G* be a *m × n* genotype matrix from *n* individuals at *m* SNPs. The genotype vector for individual *j* is a vector of length *m* denoted by *g_j_ ∈* [0, 2]*^m^* where each entry *g_i,j_ ∈* [0, 2] denotes the number of minor alleles carried by individual *j* at SNP *i* or the genotype dosage. Let *A* denote the matrix of standardized genotypes where each row *a_i_* has approximately mean 0 and variance 1 for SNPs in Hardy-Weinberg equilibrium.

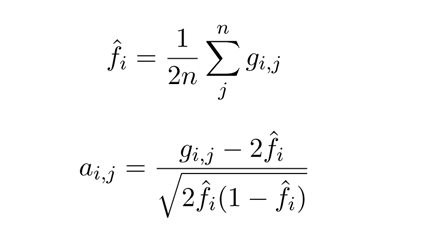

### Normalization for single-cell RNA sequencing data

PCAone supports a general CSV format as input where each row indicates the samples for the feature. By default, PCAone assumes the data in CSV format is already normalized so that users can apply their own normalization methods. We also implemented the count per median (CPMED) normalization method for scRNA-seq data in PCAone (with option *–cpmed*), which is the commonly used one (Shekhar et al. 2016; van Dijk et al. 2018) and the one implemented by OnlinePCA.jl (Tsuyuzaki et al. 2020). Therefore, we can benchmark the performance of PCAone and OnlinePCA.jl with the same normalization method. Let *X* be the raw matrix with *m* genes and *n* cells where each row *x_i_* is a vector of length *n* and each entry *x_i,j_*denotes the read counts or TPM for cell *i* at gene *j*. Let *s* = [*s*_1_*, s*_2_*, …, s_n_*] be a vector where each entry is sum of each column of *X*. Let *A* denote the normalized matrix where each entry *a_i,j_* is median-normalized and log-transformed as the follows.

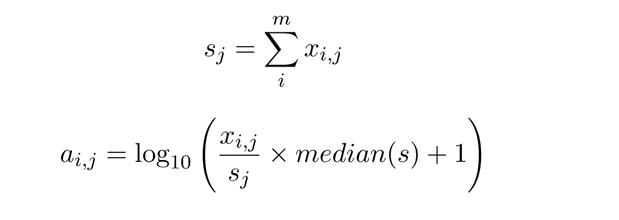

## Software availability

The source code of PCAone can be found in the Supplemental Code. Also, the PCAone C++ program is available at https://github.com/Zilong-Li/PCAone. The R package of PCAone algorithm is on CRAN https://CRAN.R-project.org/package=pcaone. The reproducible work-flow used to perform the analyses in the paper is available at https://github.com/Zilong-Li/ Papers/tree/main/pcaone.

## Acknowledgements

The study was supported by the Lundbeck foundation (R215-2015-4174) and the Novo Nordisk Foundation (NNF20OC0061343). This research has been conducted using the UK Biobank Resource under Application Number 32683. ZL and AA conceived the study and derived the methods with input from JM. ZL implemented the methods and performed the analyses. ZL, JM and AA discussed the results and contributed to writing the manuscript.

## References

1. Abraham, G, Y Qiu, and M Inouye (2017). “FlashPCA2: principal component analysis of Biobank-scale genotype datasets”. In: Bioinformatics (Oxford, England) 33.17, pp. 2776–2778. doi: 10.1093/bioinformatics/btx299.

2. Agrawal, A, AM Chiu, M Le, E Halperin, and S Sankararaman (2020). “Scalable probabilistic PCA for large-scale genetic variation data”. In: PLOS Genetics 16.5, e1008773. doi: 10.1371/journal.pgen.1008773.

3. Bansal, V, A Bashir, and V Bafna (2007). “Evidence for large inversion polymorphisms in the human genome from HapMap data”. In: Genome Research 17.2, pp. 219–230. doi: 10.1101/gr.5774507.

4. Beleza, S, AM Santos, B McEvoy, I Alves, C Martinho, E Cameron, MD Shriver, EJ Parra, and J Rocha (2013). “The Timing of Pigmentation Lightening in Europeans”. In: Molecular Biology and Evolution 30.1, pp. 24–35. doi: 10.1093/molbev/mss207.

5. Bose, A, V Kalantzis, EM Kontopoulou, M Elkady, P Paschou, and P Drineas (2019). “TeraPCA: a fast and scalable software package to study genetic variation in tera-scale genotypes”. In: Bioinformatics 35.19, pp. 3679–3683. doi: 10.1093/bioinformatics/btz157.

6. Cappé, O and E Moulines (2009). “On-line expectation–maximization algorithm for latent data models”. In: Journal of the Royal Statistical Society: Series B (Statistical Methodology) 71.3, pp. 593–613. doi: 10.1111/j.1467-9868.2009.00698.x.

7. Chang, CC, CC Chow, LC Tellier, S Vattikuti, SM Purcell, and JJ Lee (2015). “Second-generation PLINK: rising to the challenge of larger and richer datasets”. In: GigaScience 4.1, p. 7. doi: 10.1186/s13742-015-0047-8.

8. Diaz-Papkovich, A, L Anderson-Trocmé, C Ben-Eghan, and S Gravel (2019). “UMAP reveals cryptic population structure and phenotype heterogeneity in large genomic cohorts”. In: PLOS Genetics 15.11, e1008432. doi: 10.1371/journal.pgen.1008432.

9. Elgamal, T and M Hefeeda (2015). Analysis of PCA Algorithms in Distributed Environments. doi: 10.48550/arXiv.1503.05214. arXiv: 1503.05214 [cs].

10. Galinsky, KJ, G Bhatia, PR Loh, S Georgiev, S Mukherjee, NJ Patterson, and AL Price (2016). “Fast Principal-Component Analysis Reveals Convergent Evolution of ADH1B in Europe and East Asia”. In: The American Journal of Human Genetics 98.3, pp. 456–472. doi: 10.1016/j.ajhg.2015.12.022.

11. Halko, N, PG Martinsson, and JA Tropp (2011). “Finding Structure with Randomness: Probabilistic Algorithms for Constructing Approximate Matrix Decompositions”. In: SIAM Review 53.2, pp. 217–288. doi: 10.1137/090771806.

12. Kane, MJ, J Emerson, and S Weston (2013). “Scalable Strategies for Computing with Massive Data”. In: Journal of Statistical Software 55.14, pp. 1–19.

13. Kingma, DP and J Ba (2017). Adam: A Method for Stochastic Optimization. doi: 10.48550/arXiv.1412.6980. arXiv: 1412.6980 [cs].

14. Kiselev, VY, TS Andrews, and M Hemberg (2019). “Challenges in unsupervised clustering of single-cell RNA-seq data”. In: Nature Reviews Genetics 20.5, pp. 273–282. doi: 10.1038/s41576-018-0088-9.

15. Kuhn, E, C Matias, and T Rebafka (2020). “Properties of the stochastic approximation EM al-gorithm with mini-batch sampling”. In: Statistics and Computing 30.6, pp. 1725–1739. doi: 10.1007/s11222-020-09968-0.

16. Lehoucq, RB and DC Sorensen (1996). “Deflation Techniques for an Implicitly Restarted Arnoldi Iteration”. In: SIAM Journal on Matrix Analysis and Applications 17.4, pp. 789–821. doi: 10.1137/S0895479895281484.

17. Lehoucq, RB, DC Sorensen, and C Yang (1998). ARPACK Users’ Guide: Solution of Large-Scale Eigenvalue Problems with Implicitly Restarted Arnoldi Methods. Society for Industrial et al. doi: 10.1137/1.9780898719628.

18. Levey, A and M Lindenbaum (2000). “Sequential Karhunen-Loeve basis extraction and its ap-plication to images”. In: IEEE Transactions on Image Processing 9.8, pp. 1371–1374. doi: 10.1109/83.855432.

19. Maaten, L van der and G Hinton (2008). “Visualizing Data using t-SNE”. In: Journal of Machine Learning Research 9.86, pp. 2579–2605.

20. Meisner, J and A Albrechtsen (2018). “Inferring Population Structure and Admixture Proportions in Low-Depth NGS Data”. In: Genetics 210.2, pp. 719–731. doi: 10.1534/genetics.118. 301336.

21. Meisner, J, S Liu, M Huang, and A Albrechtsen (2021). “Large-scale inference of population structure in presence of missingness using PCA”. In: Bioinformatics 37.13, pp. 1868–1875. doi: 10.1093/bioinformatics/btab027.

22. Neal, RM and GE Hinton (1998). “A View of the Em Algorithm that Justifies Incremental, Sparse, and other Variants”. In: Learning in Graphical Models. Ed. by MI Jordan. NATO ASI Series. Dordrecht: Springer Netherlands, pp. 355–368. doi: 10.1007/978-94-011-5014-9_12.

23. Patterson, NJ, AL Price, and D Reich (2006). “Population Structure and Eigenanalysis”. In: PLOS Genetics 2.12, e190. doi: 10.1371/journal.pgen.0020190.

24. Porubsky, D et al. (2022). “Recurrent inversion polymorphisms in humans associate with genetic instability and genomic disorders”. In: Cell 0.0. doi: 10.1016/j.cell.2022.04.017.

25. Price, AL, NJ Patterson, RM Plenge, ME Weinblatt, NA Shadick, and D Reich (2006). “Principal components analysis corrects for stratification in genome-wide association studies”. In: Nature Genetics 38.8, pp. 904–909. doi: 10.1038/ng1847.

26. Privé, F, H Aschard, A Ziyatdinov, and MGB Blum (2018). “Efficient analysis of large-scale genome-wide data with two R packages: bigstatsr and bigsnpr”. In: Bioinformatics 34.16, pp. 2781–2787. doi: 10.1093/bioinformatics/bty185.

27. Privé, F, K Luu, MGB Blum, JJ McGrath, and BJ Vilhjálmsson (2020). “Efficient toolkit implementing best practices for principal component analysis of population genetic data”. In: Bioinformatics 36.16, pp. 4449–4457. doi: 10.1093/bioinformatics/btaa520.

28. R Core Team (n.d.). R: A Language and Environment for Statistical Computing. R Foundation for Statistical Computing. Vienna, Austria.

29. Ross, DA, J Lim, RS Lin, and MH Yang (2008). “Incremental Learning for Robust Visual Tracking”. In: International Journal of Computer Vision 77.1–3, pp. 125–141. doi: 10.1007/s11263-007-0075-7.

30. Shekhar, K et al. (2016). “Comprehensive Classification of Retinal Bipolar Neurons by Single-Cell Transcriptomics”. In: Cell 166.5, 1308–1323.e30. doi: 10.1016/j.cell.2016.07.054.

31. Tsuyuzaki, K, H Sato, K Sato, and I Nikaido (2020). “Benchmarking principal component analysis for large-scale single-cell RNA-sequencing”. In: Genome Biology 21.1, p. 9. doi: 10.1186/s13059-019-1900-3.

32. van Dijk, D et al. (2018). “Recovering Gene Interactions from Single-Cell Data Using Data Diffusion”. In: Cell 174.3, 716–729.e27. doi: 10.1016/j.cell.2018.05.061.

33. Yu, W, Y Gu, J Li, S Liu, and Y Li (2017). “Single-Pass PCA of Large High-Dimensional Data”. In: pp. 3350–3356.

34. Zheng, GXY et al. (2017). “Massively parallel digital transcriptional profiling of single cells”. In: Nature Communications 8, p. 14049. doi: 10.1038/ncomms14049.

