## Supplemental_Material for "Fast and accurate out-of-core PCA framework for large scale biobank data"

### Supplementary figures

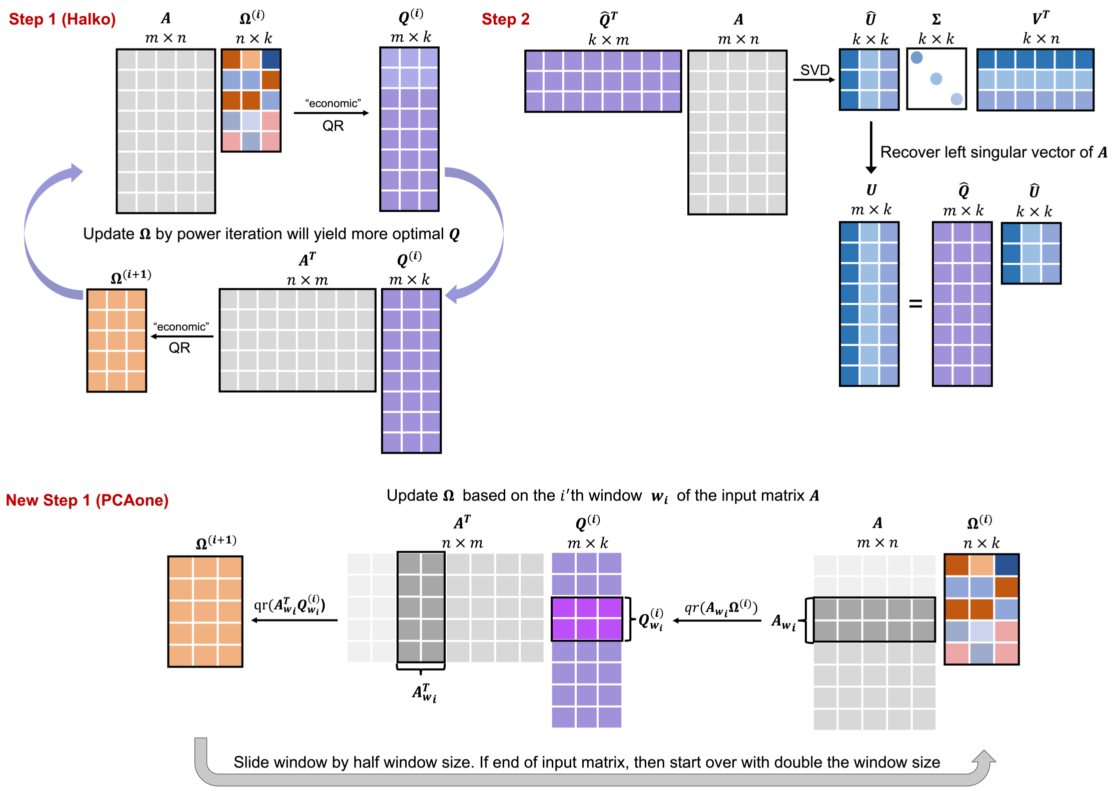


Figure S1: Overview of PCAone algorithm. The underlying idea of Halko randomized SVD is seeking a near-optimal orthonormal basis $Q^{m \times k}$ for input matrix $A^{m \times n}$, such that $A \approx QQ^T A$ is satisfied with $k \ll m, n$. In **step1 of the Halko**, $Q$ is obtained from QR orthogonalization $Q = qr(A\Omega)$ on a random low-dimensional matrix $\Omega^{n \times k}$ multiplied with the input matrix. Better accuracy is achived by iterative updating $Q$ and $\Omega$, which composes a so-called power iteration step. In **Step2**, with such a $Q$, the right eigenvectors of $A$ are approximated by the right eigenvectors of the low dimensional matrix $B = Q^T A$. For large datasets, a **new step1 in PCAone** is proposed, which uses a subset of the input matrix $A_{w_i}$ each time to perform power iterations in a sliding window with step size of half the window. After each pass through the whole data, the window size of $A_{w_i}$ is doubled until $A_{w_i}$ equals $A$.

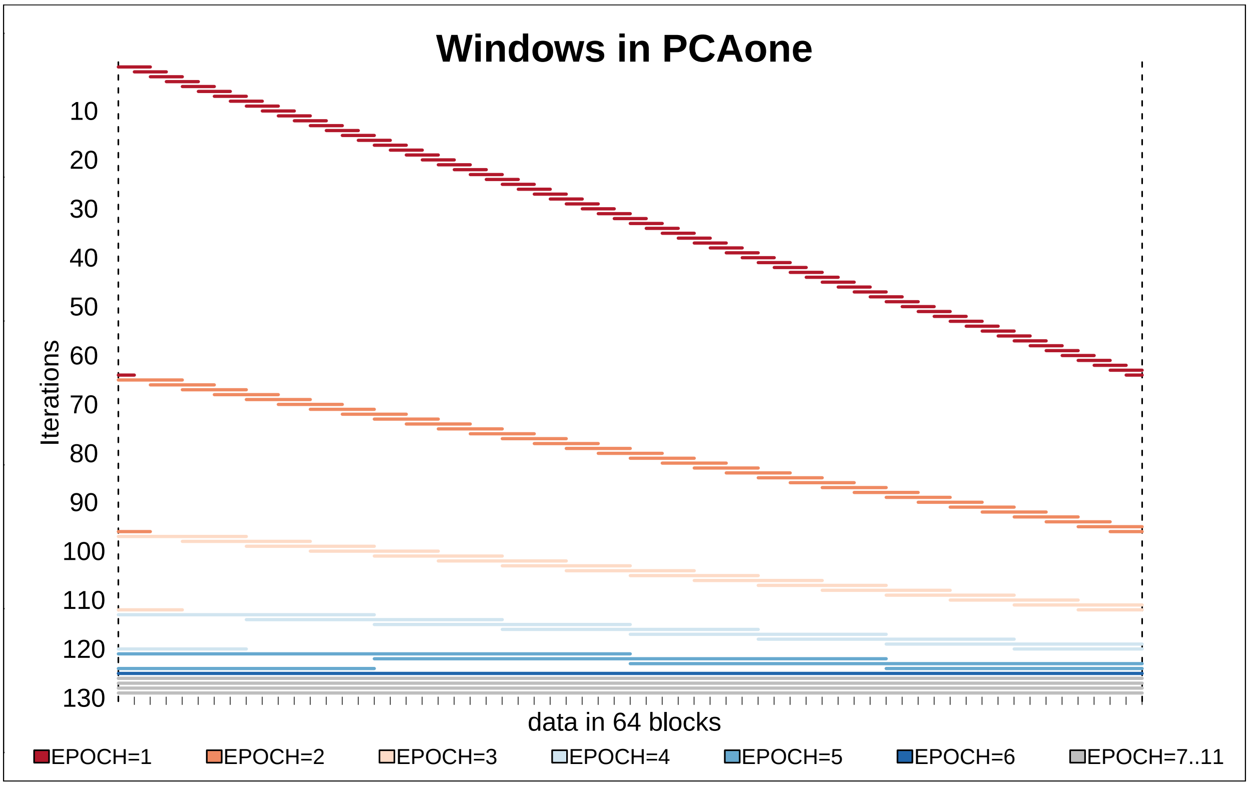


Figure S2: Illustration of the windows of data used for each iteration, where each horizontal line represents blocks of the data that is used to update $\Omega$ at each iteration. Each epoch denotes one round of reading the data from the disk and each iteration is colored by its epoch. In this illustration, the data is split into 64 blocks and in the first epoch a window size of two blocks and a step size of one block is used. The window wraps around the data such as the window starting at the last block will use data from the first one block as well. In each of the following epochs, both the window size and the step size is doubled. For the first 6 epochs the data is read 6 times from the disk but $\Omega$ is updated $64 + 32 + 16 + 8 + 4 + 1 = 124$ times where the last iteration uses the whole data.

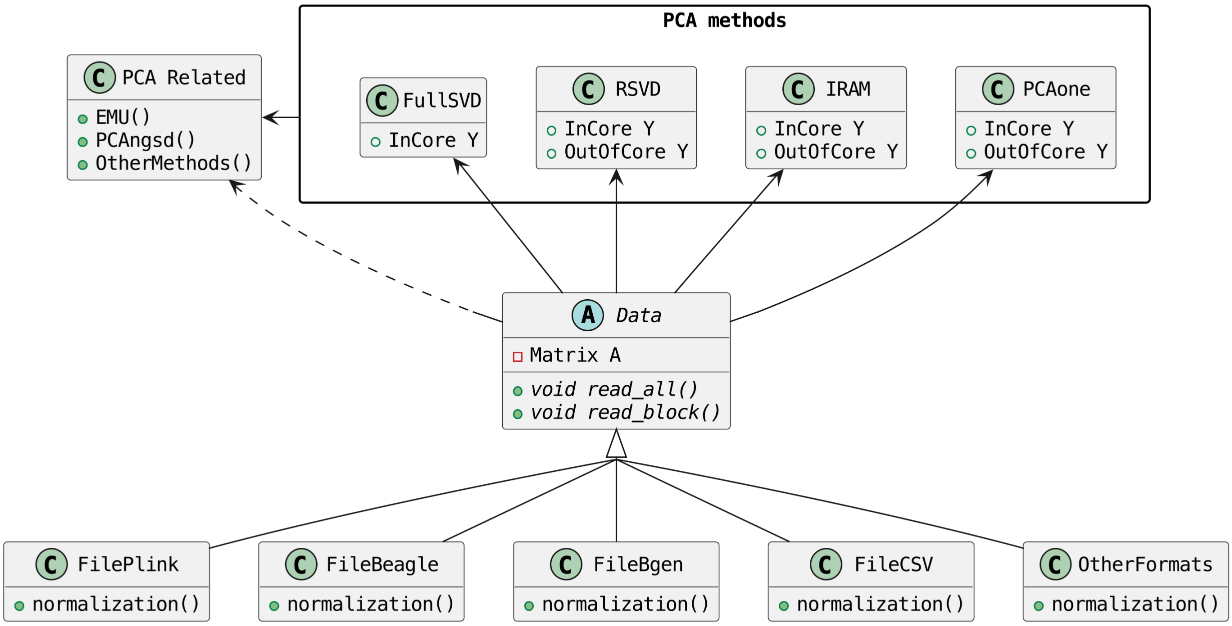


Figure S3: Overview of PCAone software architecture. There are 3 different partial SVD algorithms (IRAM, RSVD, PCAone) with both in-core and out-of-core implementations and one full SVD in-core implementation in PCAone software. Multiple file formats are supported, i.e. FilePlink for diallelic genotype data (PLINK), FileBeagle for genotype likelihood data (BEAGLE), FileBgen for allelic dosages from imputation (BGEN), and FileCSV with real values for other data type (CSV), such as single cell and bulk RNA-seq data. PCAone also works for genotype data with missingness using EMU algorithm and genotype likelihoods from low depth sequencing data using PCAngsd algorithm.

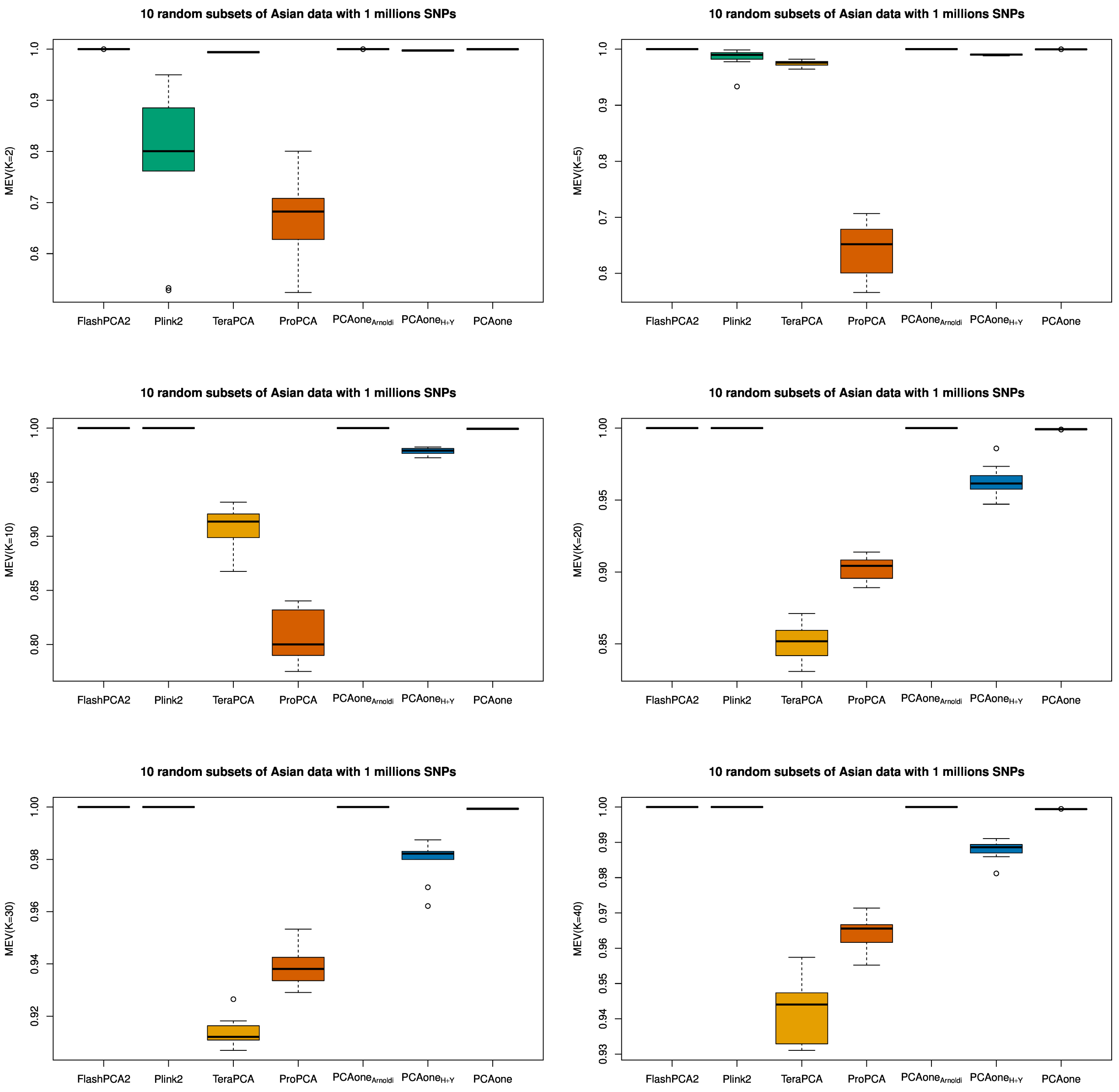


Figure S4: MEV uncertainty of different methods for different estimated PCs. We randomly subset one million SNPs of the East Asian data for 10 times to access the uncertainty of MEV estimates. As shown, the MEV estimates are consistent for all methods which confirms conclusions in the main text.

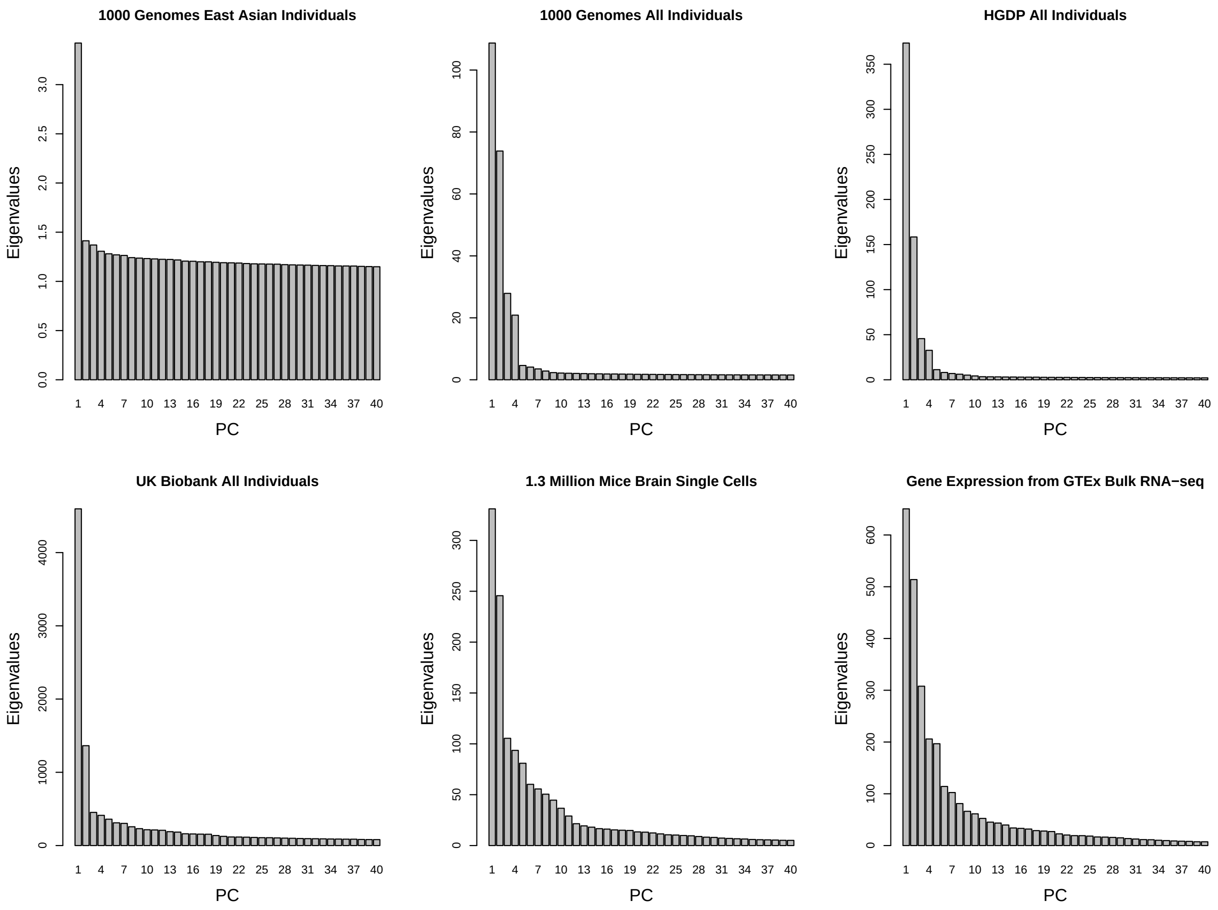


Figure S5: Top 40 eigenvalues for different datasets. For the 1000 Genomes, East Asian and HGDP, the results are from full SVD. For UK Biobank, single cell RNA-seq and bulk RNA-seq, the results are from $\text{PCAone}_{Arnoldi}$

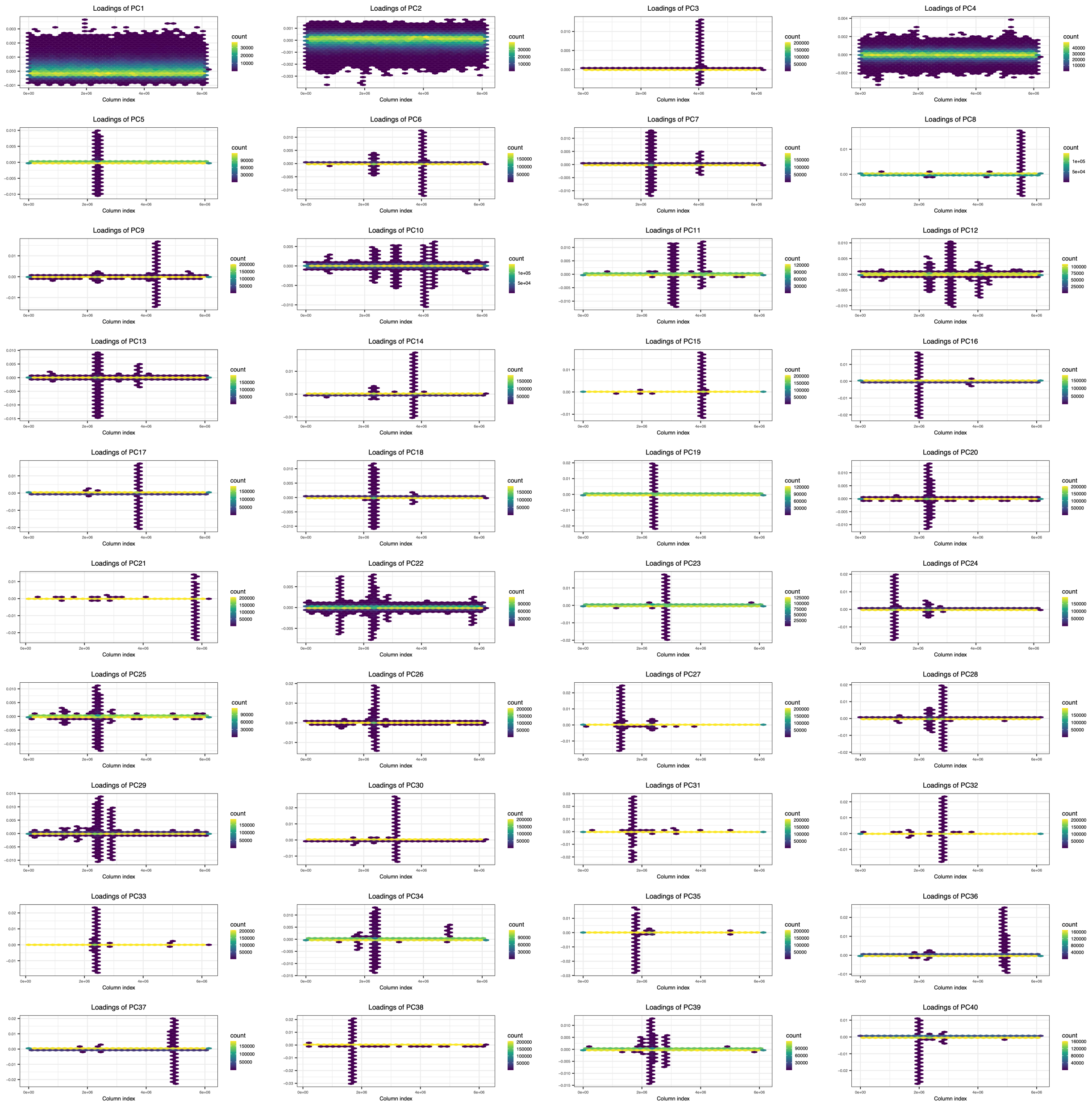


Figure S6: SNP loadings of top 40 PCs for the UK Biobank data with 487,409 individuals and 6,133,304 SNPs. SNPs are ordered by chromosome and physical position, represented on the x-axis by their order, and the value of loadings is represented on the y-axis. Points are hex-binned and colored by their density.

**A** Number of passes over data (epochs) of each method for top K PCs

| K(PCs) | PCAone | PCAone$_{H+Y}$ | TeraPCA | Plink2(FastPCA)* | ProPCA* | FlashPCA2 | PCAone$_{Arnoldi}$ |
|---|---|---|---|---|---|---|---|
| 2 | 2 | 3 | 6 | 2 | 4 | 10 | 21 |
| 5 | 4 | 6 | 12 | 5 | 7 | 25 | 21 |
| 10 | 7 | 11 | 22 | 10 | 12 | 55 | 46 |
| 20 | 7 | 18 | 42 | 20 | 22 | 117 | 98 |
| 30 | 7 | 19 | 30 | 30 | 32 | 172 | 144 |
| 40 | 7 | 19 | 36 | 40 | 42 | 223 | 188 |

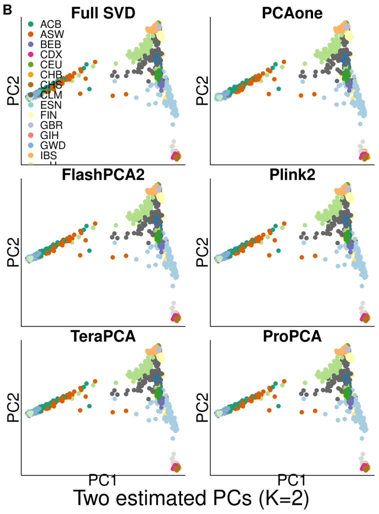


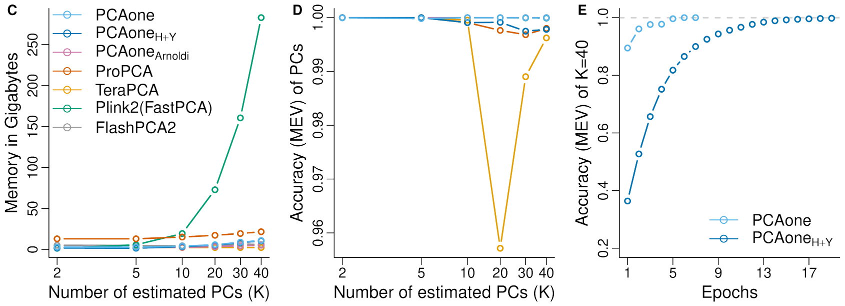


Figure S7: Performance on 1000 Genomes Project data with all populations. PCA performance of different software by varying the number of inferred PCs (K) based on 5,766,022 SNPs and 2,504 individuals. **(A)**, Number of epochs used by each software. $*$ Not out-of-core, only allows for in-core computation with this number of iterations. **(B)**, Two estimated PCs $K = 2$ for each methods including results of the full-rank SVD. **(C)**, Memory usage as a function of $K$. **(D)**, Accuracy (MEV) compared to the full-rank SVD as a function of $K$. **(E)**, Convergence of PCAone and PCAone$_{H+Y}$ shown as accuracy per epoch.

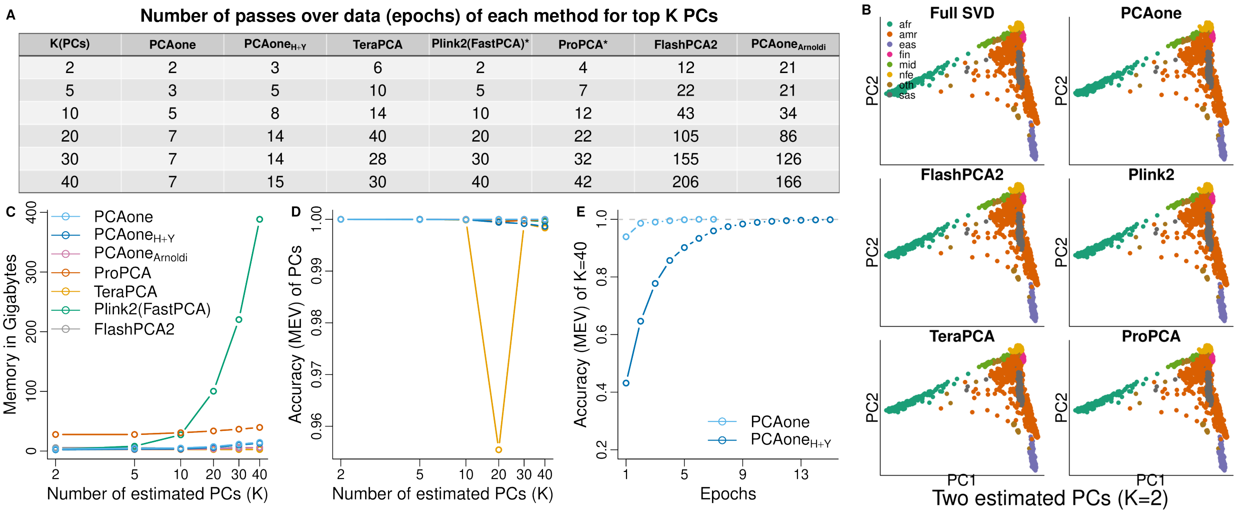


| K(PCs) | PCAone | $\text{PCAone}_{H+Y}$ | TeraPCA | Plink2(FastPCA)* | ProPCA* | FlashPCA2 | $\text{PCAone}_{Arnoldi}$ |
|---|---|---|---|---|---|---|---|
| 2 | 2 | 3 | 6 | 2 | 4 | 12 | 21 |
| 5 | 3 | 5 | 10 | 5 | 7 | 22 | 21 |
| 10 | 5 | 8 | 14 | 10 | 12 | 43 | 34 |
| 20 | 7 | 14 | 40 | 20 | 22 | 105 | 86 |
| 30 | 7 | 14 | 28 | 30 | 32 | 155 | 126 |
| 40 | 7 | 15 | 30 | 40 | 42 | 206 | 166 |

Figure S8: Performance on HGDP data with all populations. PCA performance of different software by varying the number of inferred PCs (K) based on 7,915,146 SNPs and 4151 individuals. **(A)**, Number of epochs used by each software. $*$ Not out-of-core, only allows for in-core computation with this number of iterations. **(B)**, Two estimated PCs $K = 2$ for each methods including results of the full-rank SVD. **(C)**, Memory usage as a function of $K$. **(D)**, Accuracy (MEV) compared to the full-rank SVD as a function of $K$. **(E)**, Convergence of PCAone and $\text{PCAone}_{H+Y}$ shown as accuracy per epoch.

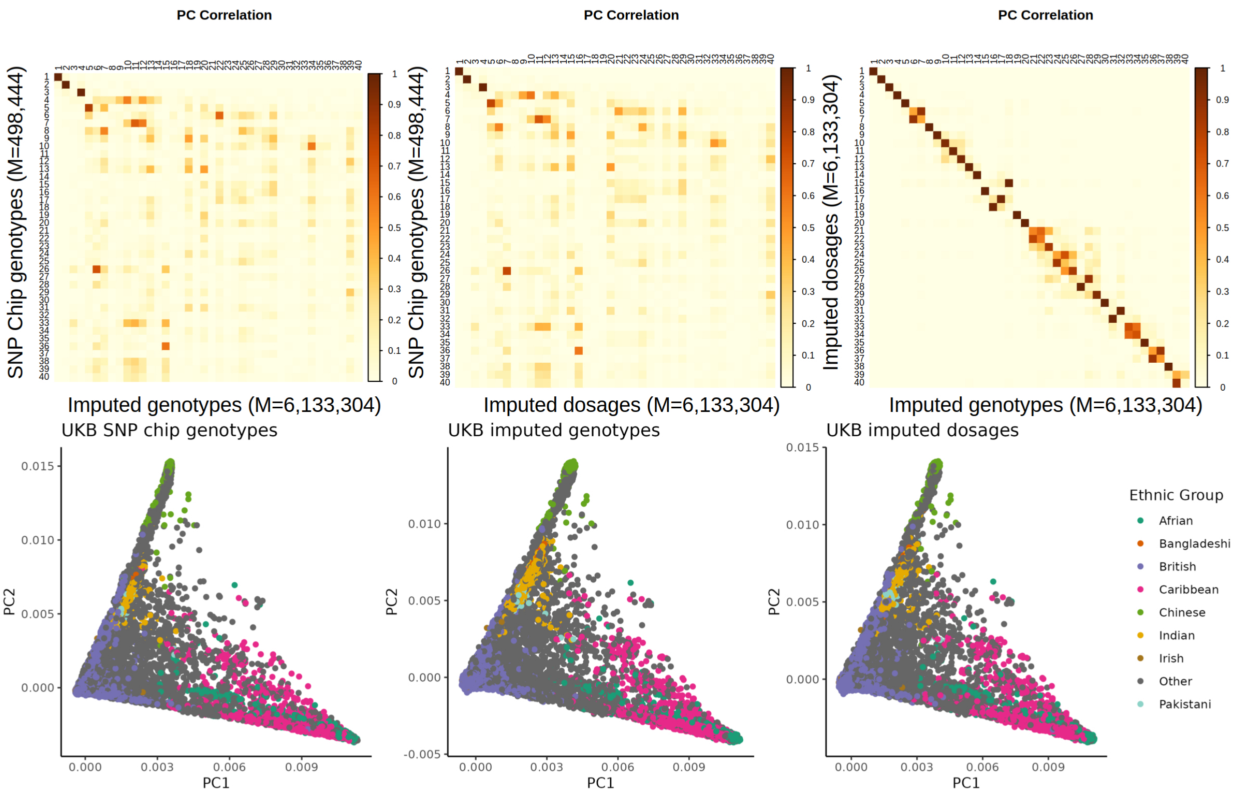


Figure S9: PCA analyses on UK Biobank SNP chip genotypes, imputed genotypes and imputed genotypes. As shown in the main, PC1, PC2 and PC4 capture population structure. And there is also high correlation between the imputed genotypes (6,133,304 SNPs) and SNP chip genotypes (498,444 SNPs) for PC1, PC2 and PC4. The correlations for most PCs between the imputed genotypes and dosages are close to 1.0. The speed performance of each dataset in different formats is shown in Table S3

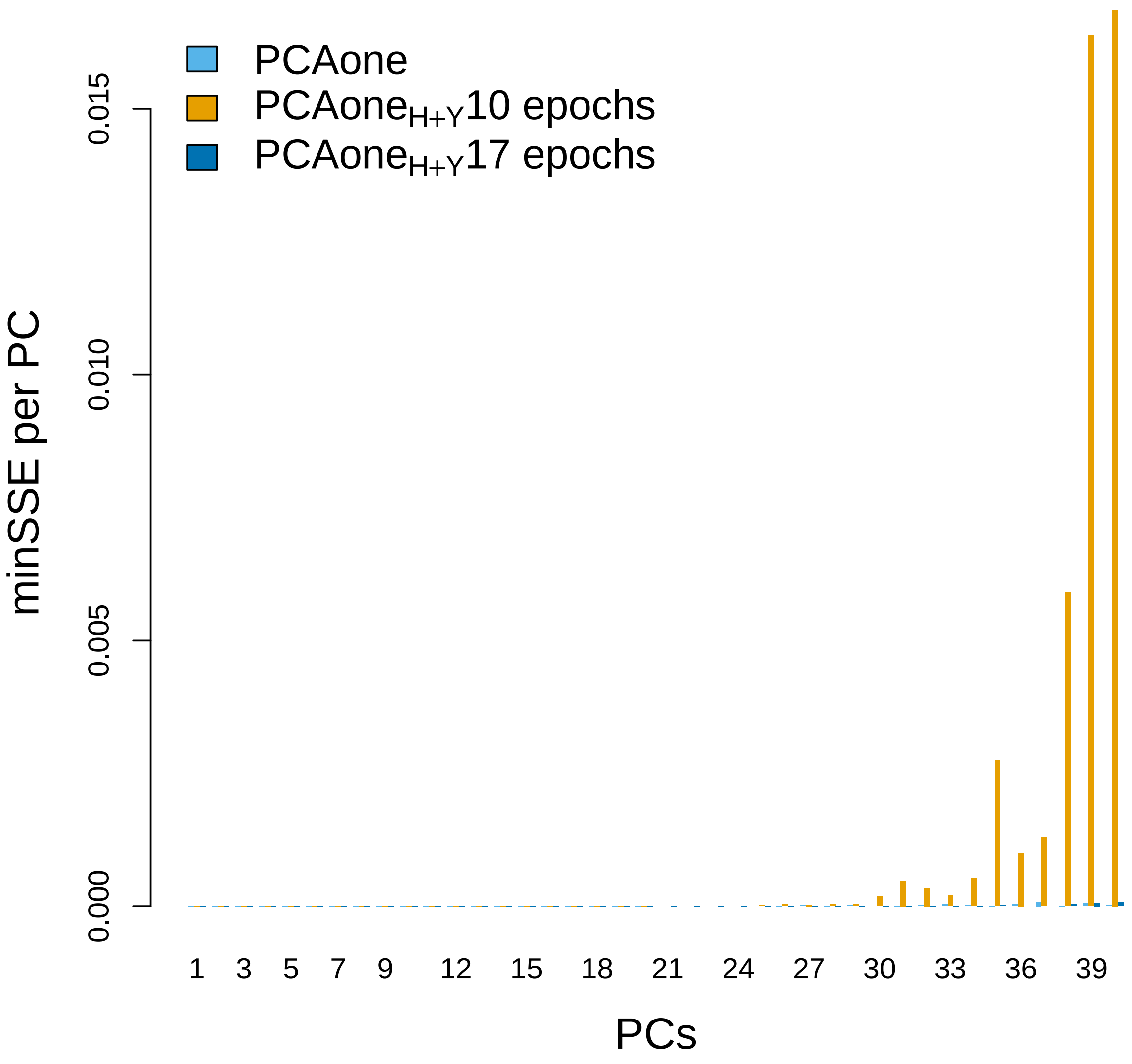


Figure S10: minSSE of top 40 PCs of the UK Biobank data with 487,409 individuals and 6,133,304 SNPs. minSSE were calculated based on each PC with the lowest distance to the PCs from PCAone$_{Arnoldi}$.

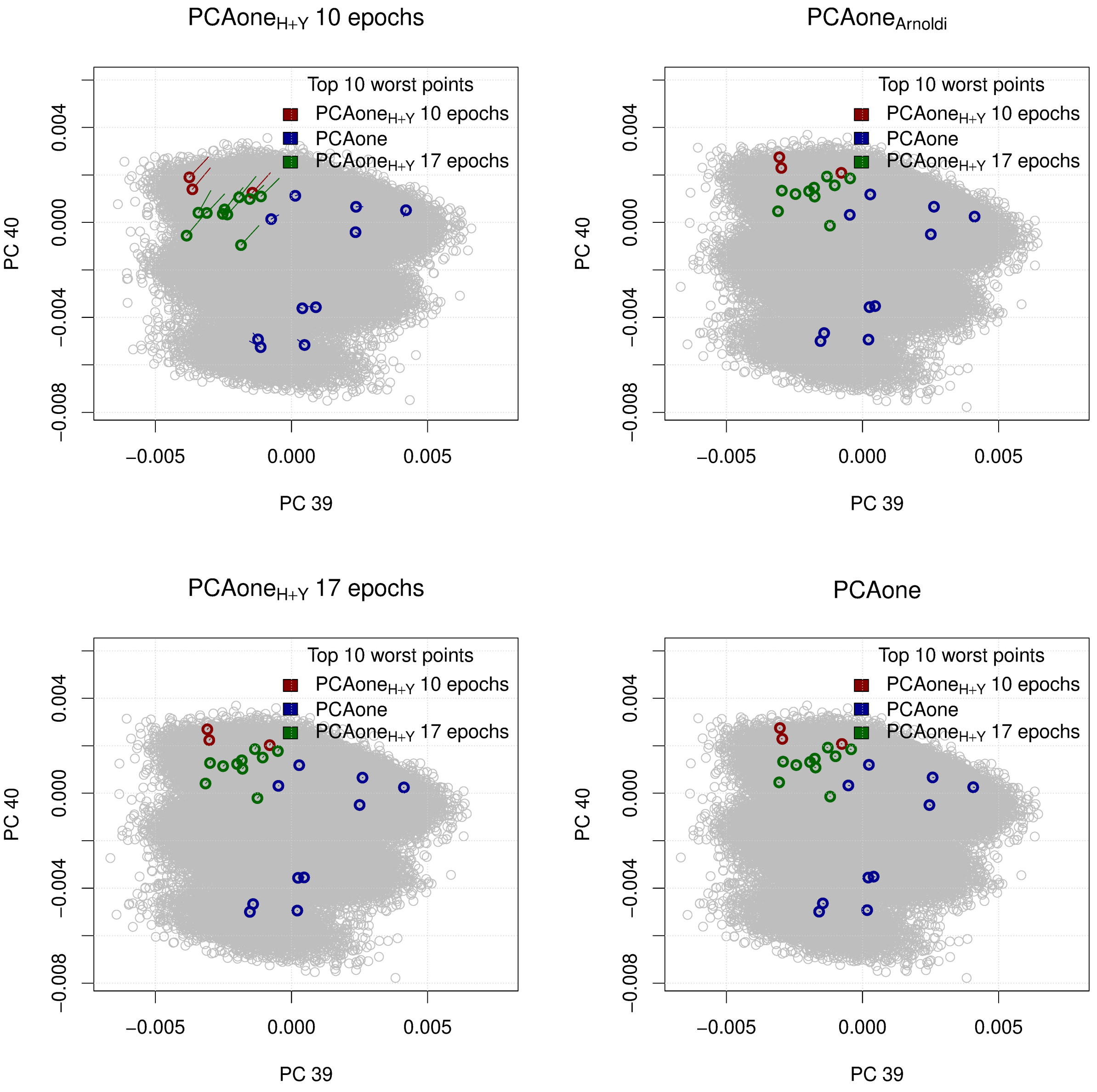


Figure S11: PC 39 and 40 of the UK Biobank data with 487,409 individuals and 6,133,304 SNPs. The top 10 points with the worst distance (minSSE) to the results from PCAone$_{Arnoldi}$ are highlighted , and each point has a line that starts in the position in the PCAone$_{Arnoldi}$ coordinates.

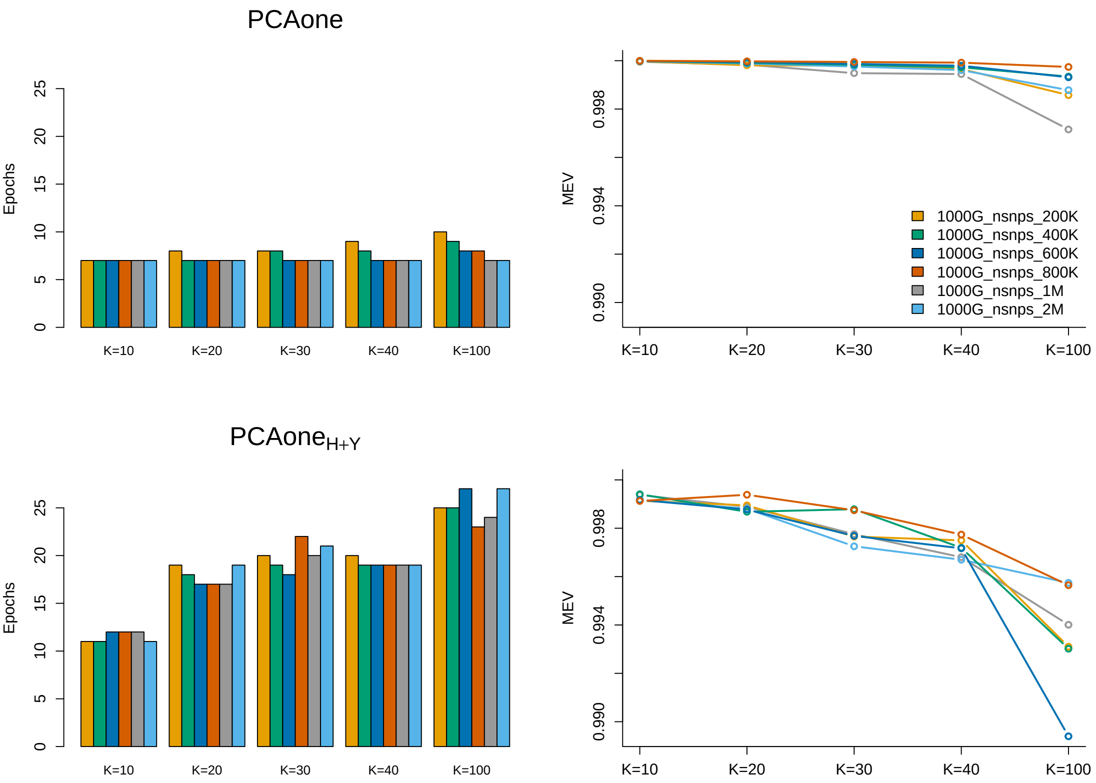


(a) 1000 genomes dataset with N=2,504 samples

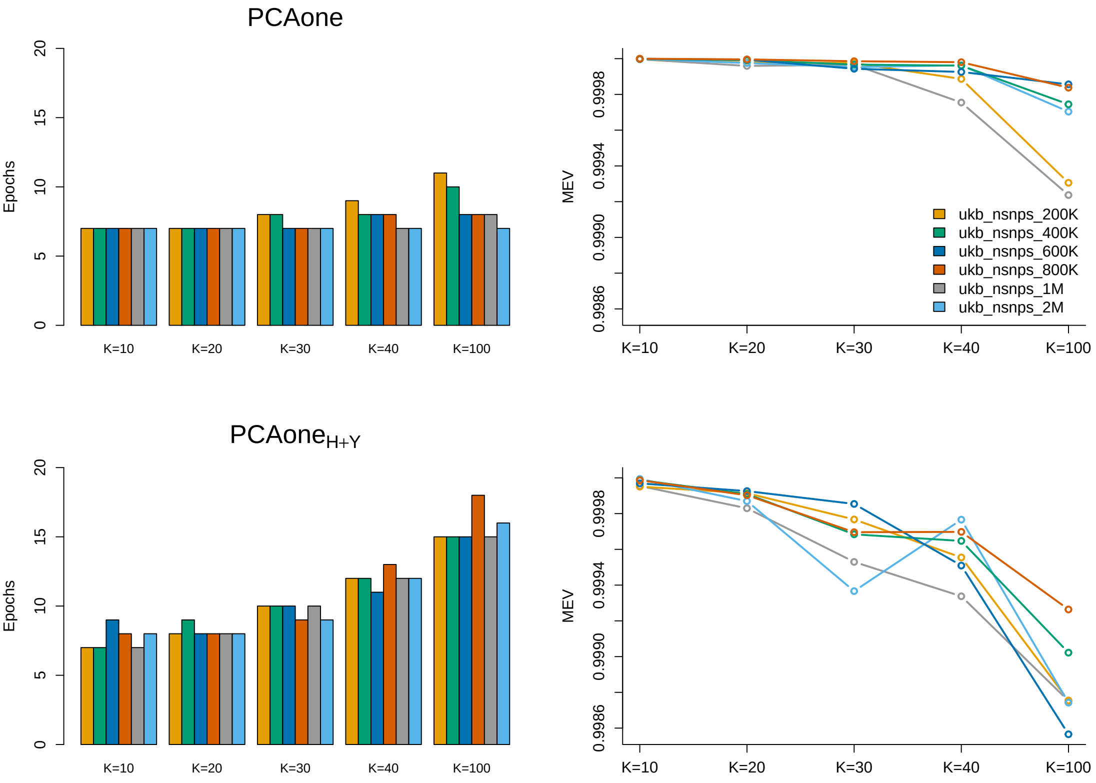


(b) UK Biobank dataset with random N=50,000 samples

Figure S12: PCAone performance on dataset scaled by the number of features with fixed number of samples. **(a)**, Datasets are randomly subset from the 1000 genomes data with 2,504 samples. **(b)**, Datasets are randomly subset from the UK Biobank data with 50,000 samples.

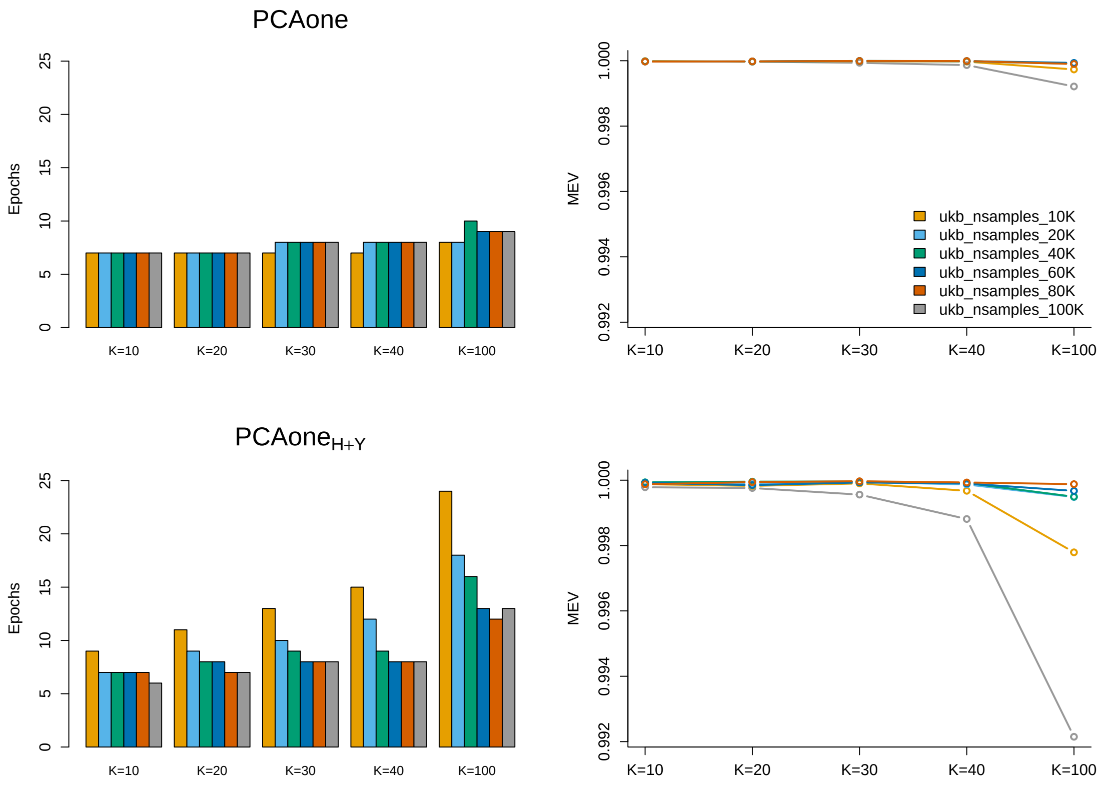


(a) UK Biobank array dataset with M=671,191 SNPs

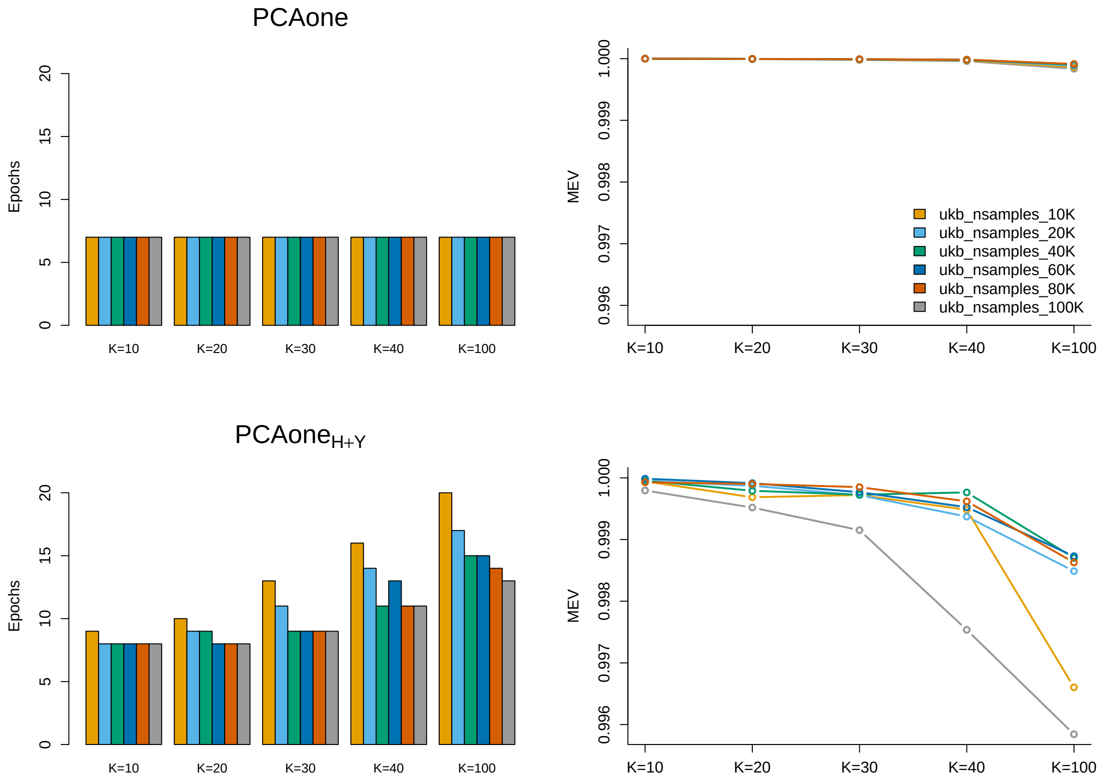


(b) UK Biobank imputed dataset with random M=3,000,000 SNPs

Figure S13: PCAone performance on dataset scaled by the number of samples with fixed number of features. **(a)**, Datasets are randomly subset from the UK Biobank SNP chip data with 671,191 SNPs. **(b)**, Datasets are randomly subset from the UK Biobank imputed data with 3,000,000 SNPs.

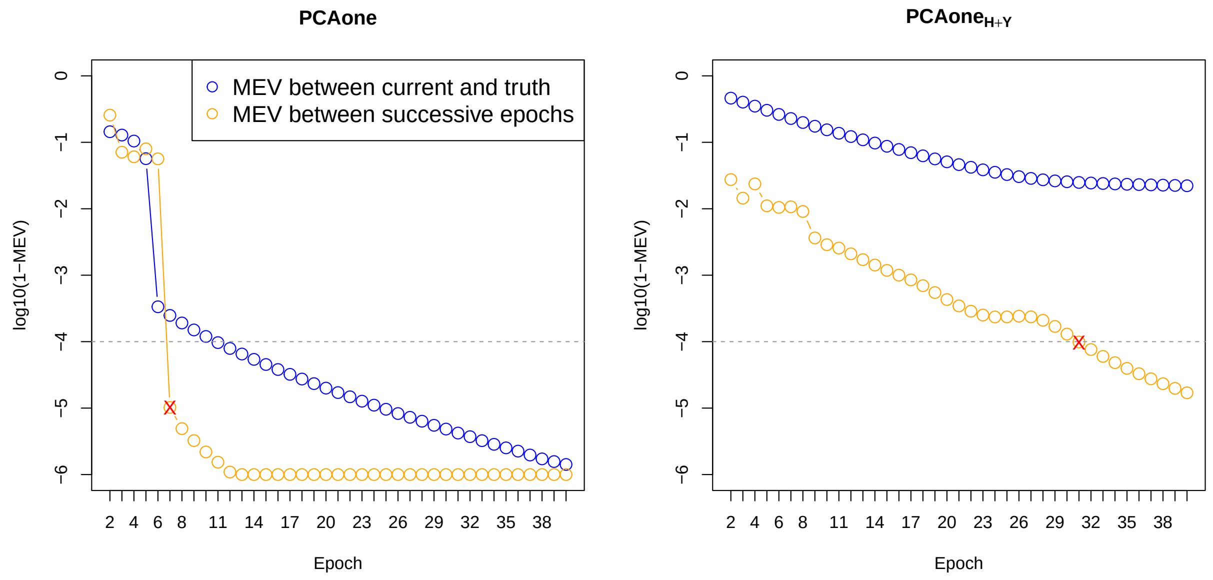


(a) Asian dataset

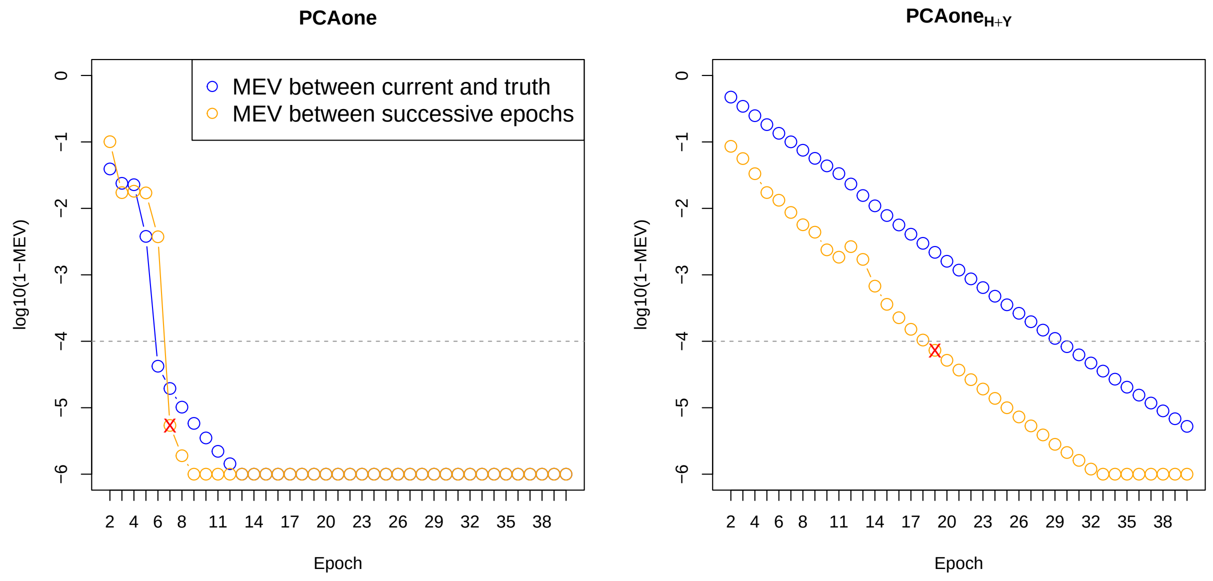


(b) 1000 genomes dataset

Figure S14: Distribution for the estimated MEV of top $K = 40$ PCs between successive epochs and the true MEV of PCs from each epoch against full SVD. It's shown that the estimated MEV (orange line) follows the same trend as the true MEV (blue line). Hence the more epochs PCAone runs, the more accurate the results would be. We use a cut-off value of $1 - MEV < 10^{-4}$ (gray line) as the stopping criteria in default for two RSVD methods (PCAone and PCAone$_{H+Y}$), which gives a good balance between the accuracy and epochs. As shown, PCAone can achieve a true MEV larger than 0.9999 for both datasets while using only 7 epochs.

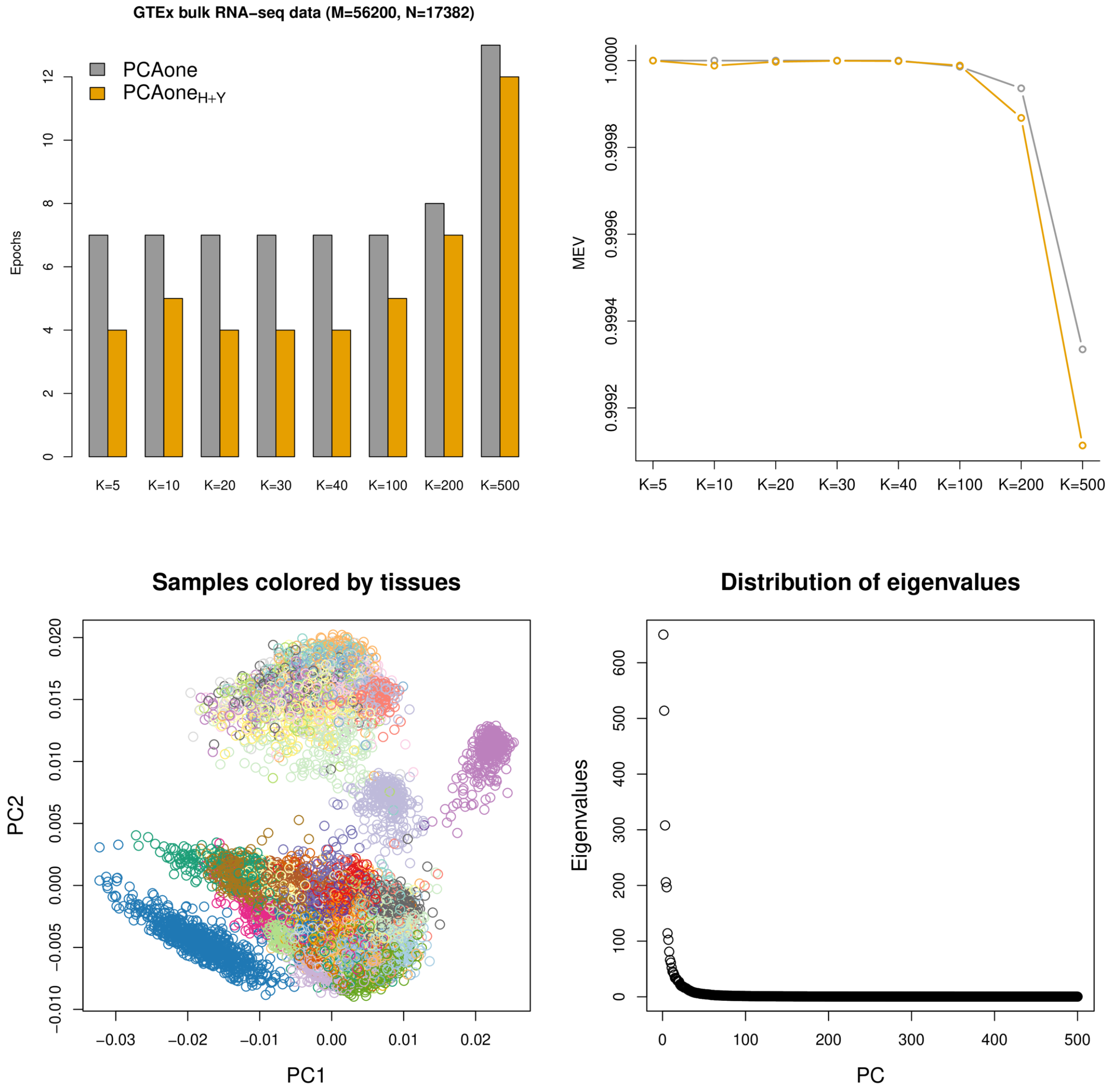


Figure S15: Analyses on bulk RNA sequencing data from the GTEx project, consisting of M=56,200 transcript counts (TPM) and N=17,382 samples from 54 tissues and 943 individuals. PCA results by PCAone$_{Arnoldi}$ are used for calculating MEV and displaying eigenvalues. The points in PCA plot are colored by the tissue catagory.

### Supplementary tables

Table S1: MEV of top 40 PCs for 13 random subsets of UK Biobank imputed data in Figure 2. (*) As it is not computationally feasible to perform the full SVD for large datasets, we therefore use PCAone$_{Arnoldi}$ as the truth to calculate the accuracy, which is indicated by "-". Some analysis did not finish due to being out of memory (oom, 180G) and out of time(oot, 1 day).

| N | M | PCAone | PCAone$_{H+Y}$ | Plink2 | TeraPCA | ProPCA | FlashPCA2 | PCAone$_{Arnoldi}$ * |
|---|---|---|---|---|---|---|---|---|
| 10K | 3M | 1.0000 | 0.9984 | oom | 0.9978 | 0.9997 | 1.0000 | - |
| 20K | 3M | 1.0000 | 0.9993 | oom | 0.9980 | 1.0000 | 1.0000 | - |
| 40K | 3M | 1.0000 | 0.9995 | oom | 0.9995 | 1.0000 | 1.0000 | - |
| 60K | 3M | 1.0000 | 0.9996 | oom | 0.9995 | 1.0000 | 1.0000 | - |
| 80K | 3M | 1.0000 | 0.9996 | oom | 0.9995 | 1.0000 | oot | - |
| 100K | 3M | 1.0000 | 0.9998 | oom | 0.9996 | 1.0000 | oot | - |
| 50K | 200K | 0.9997 | 0.9994 | 1.0000 | 0.9994 | 1.0000 | 1.0000 | 1.0000 |
| 50K | 400K | 1.0000 | 0.9996 | 1.0000 | 0.9989 | 1.0000 | 1.0000 | 1.0000 |
| 50K | 600K | 1.0000 | 0.9997 | 1.0000 | 0.9995 | 1.0000 | 1.0000 | 1.0000 |
| 50K | 800K | 1.0000 | 0.9993 | 1.0000 | 0.9993 | 0.9999 | 1.0000 | 1.0000 |
| 50K | 1M | 1.0000 | 0.9996 | 1.0000 | 0.9996 | 1.0000 | 1.0000 | 1.0000 |
| 50K | 2M | 1.0000 | 0.9995 | oom | 0.9992 | 1.0000 | 1.0000 | 1.0000 |
| 50K | 3M | 1.0000 | 0.9995 | oom | 0.9995 | 1.0000 | 1.0000 | 1.0000 |

Table S2: minSSE of top 40 PCs for 13 random subsets of UK Biobank imputed data in Figure 2. (*) As it is not computationally feasible to perform the full SVD for large datasets, we therefore use PCAone$_{Arnoldi}$ as the truth to calculate the accuracy, which is indicated by "-". Some analysis did not finish due to being out of memory (oom, 180G) and out of time(oot, 1 day).

| N | M | PCAone | PCAone$_{H+Y}$ | Plink2 | TeraPCA | ProPCA | FlashPCA2 | PCAone$_{Arnoldi}$ * |
|---|---|---|---|---|---|---|---|---|
| 10K | 3M | 0.0014 | 0.2695 | oom | 0.2086 | 0.0266 | 1e-10 | - |
| 20K | 3M | 0.0016 | 0.0630 | oom | 0.1875 | 0.0020 | 1e-10 | - |
| 40K | 3M | 0.0014 | 0.0706 | oom | 0.0491 | 1e-4 | 1e-10 | - |
| 60K | 3M | 0.0021 | 0.1541 | oom | 0.0817 | 3e-4 | 1e-10 | - |
| 80K | 3M | 0.0014 | 0.1198 | oom | 0.0418 | 6e-5 | oot | - |
| 100K | 3M | 0.0012 | 0.0558 | oom | 0.0412 | 4e-5 | oot | - |
| 50K | 200K | 0.0279 | 0.0595 | 0.0003 | 0.1091 | 0.0018 | 1e-10 | 1e-10 |
| 50K | 400K | 0.0042 | 0.0422 | 0.0002 | 0.1779 | 0.0010 | 1e-10 | 1e-10 |
| 50K | 600K | 0.0033 | 0.0348 | 0.0001 | 0.0498 | 1e-5 | 1e-10 | 1e-10 |
| 50K | 800K | 0.0051 | 0.0954 | 0.0002 | 0.2134 | 0.0070 | 1e-10 | 1e-10 |
| 50K | 1M | 0.0033 | 0.0449 | 0.0001 | 0.0746 | 1e-4 | 1e-10 | 1e-10 |
| 50K | 2M | 0.0025 | 0.0574 | oom | 0.1021 | 6e-4 | 1e-10 | 1e-10 |
| 50K | 3M | 0.0013 | 0.0769 | oom | 0.0566 | 3e-4 | 1e-10 | 1e-10 |

Table S3: PCAone performance on different UK Biobank dataset for 40 PCs using 20 threads and 20 GB memory. * refers to elpased time for shuffling the input file.

| Dataset | N | M | Format | File size | Epoch | Runtime(h) |
|---|---|---|---|---|---|---|
| Imputed genotypes | 487,409 | 6,133,304 | Plink | 697G | 7 | 9.2(8.3+0.9*) |
| Imputed dosages | 487,409 | 6,133,304 | Bgen | 1.1T | 7 | 30.5(27.1+3.4*) |
| SNP chip genotypes | 487,409 | 498,444 | Plink | 57G | 8 | 0.71 |

Table S4: MEV against PCs estimated by full-rank SVD (bold) and PCAone$_{Arnoldi}$ as a function of top PCs for scRNAs data with 12000 cells and 23771 genes. The default three iterations was used for OnlinePCA$_{Halko}$.

| PCs | PCAone | | PCAone$_{H+Y}$ | | OnlinePCA$_{Halko}$ | |
|---|---|---|---|---|---|---|
| 2 | **1.0000000** | 1.0000000 | **1.0000000** | 1.0000000 | **0.9753135** | 0.9753135 |
| 3 | **1.0000000** | 1.0000000 | **1.0000000** | 1.0000000 | **0.9804843** | 0.9804843 |
| 4 | **1.0000000** | 1.0000000 | **1.0000000** | 1.0000000 | **0.9824888** | 0.9824888 |
| 5 | **1.0000000** | 1.0000000 | **1.0000000** | 1.0000000 | **0.9859905** | 0.9859905 |
| 6 | **1.0000000** | 1.0000000 | **1.0000000** | 1.0000000 | **0.9879956** | 0.9879956 |
| 7 | **1.0000000** | 1.0000000 | **1.0000000** | 1.0000000 | **0.9897004** | 0.9897004 |
| 8 | **1.0000000** | 1.0000000 | **1.0000000** | 1.0000000 | **0.9848932** | 0.9848932 |
| 9 | **1.0000000** | 1.0000000 | **1.0000000** | 1.0000000 | **0.9889352** | 0.9889352 |
| 10 | **1.0000000** | 1.0000000 | **1.0000000** | 1.0000000 | **0.9900309** | 0.9900309 |
| 11 | **1.0000000** | 1.0000000 | **1.0000000** | 1.0000000 | **0.9903505** | 0.9903505 |
| 12 | **1.0000000** | 1.0000000 | **1.0000000** | 1.0000000 | **0.9912341** | 0.9912341 |
| 13 | **1.0000000** | 1.0000000 | **1.0000000** | 1.0000000 | **0.9918151** | 0.9918151 |
| 14 | **1.0000000** | 1.0000000 | **1.0000000** | 1.0000000 | **0.9891819** | 0.9891819 |
| 15 | **1.0000000** | 1.0000000 | **0.9999999** | 0.9999999 | **0.9703988** | 0.9703988 |
| 16 | **1.0000000** | 1.0000000 | **0.9999998** | 0.9999998 | **0.9913479** | 0.9913479 |
| 17 | **0.9999999** | 0.9999999 | **0.9999996** | 0.9999996 | **0.9899419** | 0.9899419 |
| 18 | **0.9999999** | 0.9999999 | **0.9999987** | 0.9999987 | **0.9890437** | 0.9890437 |
| 19 | **0.9999997** | 0.9999997 | **0.9999976** | 0.9999976 | **0.9698647** | 0.9698647 |
| 20 | **0.9999995** | 0.9999995 | **0.9999940** | 0.9999940 | **0.9817112** | 0.9817112 |
| Truth | full SVD | PCAone$_{Arnoldi}$ | full SVD | PCAone$_{Arnoldi}$ | full SVD | PCAone$_{Arnoldi}$ |

Table S5: Stopping criterion used by different programs. For each epoch $i$, $V^{(i)}$ denotes matrix of sigularvectors, $s^{(i)}$ denotes vector of singularvalues, $p^{(i)}$ denotes likelihood. $k$ denotes number of top PCs to be estimated by the program.

| Program | Stopping criterion |
|---|---|
| PCAone | $1 - mev(V^{(i)}, V^{(i-1)}) < 1e - 4$ |
| PCAone$_{H+Y}$ | $1 - mev(V^{(i)}, V^{(i-1)}) < 1e - 4$ |
| FastPCA/Plink2 | $k$ number of iterations |
| TeraPCA | $abs((s^{(i)} - s^{(i-1)})/s^{(i)}) < 1e - 3$ |
| ProPCA | $p^{(i)} - p^{(i-1)} < 1e - 3$ |

Table S6: Algorithm's Complexity. $n$ denotes the number of samples. $m$ denotes the number of features. $b$ denotes the block size. $k$ denotes the number of top PCs to be estimated. $l = k + 10$.

| Program | Memory complexity | Time complexity per epoch |
|---|---|---|
| PCAone$_{Arnoldi}$/FlashPCA2 | $O(nb)$ | $O(nk^2)$ |
| PCAone | $O(mb + 2ml + 3nl)$ | $O(2nml)$ |
| PCAone$_{H+Y}$ | $O(mb + 2ml + 3nl)$ | $O(nml)$ |
| TeraPCA | $O(mb + 2ml)$ | $O(nml)$ |
| FastPCA/Plink2 | $O(nm + m(k + 1)l)$ | $O(nml)$ |
| ProPCA | $O(nm)$ | $O(nml/log_3(max(n, m))$ |
